# A Junction-Dependent Mechanism Drives Mammary Cell Intercalation for Ductal Elongation

**DOI:** 10.1101/2022.11.11.516046

**Authors:** Alexander Pfannenstein, Ian G. Macara

## Abstract

Mammary glands contain branched networks of ducts and alveoli that function to produce milk for offspring. While the murine luminal epithelium is organized as a cellular monolayer, it originates from multilayered structures called terminal end buds (TEB). The TEBs generate ducts of monolayered epithelial cells as they invade the fat pad, but little is known about underlying mechanisms. While apoptosis provides a plausible mechanism for cavitation of the ductal lumen, it does not account for elongation of ducts behind the TEBs. Our spatial calculations suggest that most cells in TEBs need to intercalate into the outermost luminal layer and that this migration of cells is the primary driver of cavitation and ductal elongation. To study the progression of multilayered to monolayered epithelium, we developed a quantitative cell culture assay that determines the efficiency of intercalation into an epithelial monolayer. Using this tool, we verified that loss of adherens junctions prevents stable integration of cells into monolayers, consistent with previous data in cultured cells and in primary tissue. Interestingly, tight junction (TJ) proteins also play a key role in this integration process. Although loss of the ZO-1 TJ protein in intercalating cells suppresses intercalation, loss of ZO-1 in the monolayer has the reverse effect, promoting intercalation – even though ZO-1 is not necessary for establishment of TJs. ZO-1-positive puncta form between cells and the monolayer, which then resolves into a new intercellular boundary as intercalation proceeds. ZO-1 loss also reduces engraftment when cells are transplanted into the mammary gland via intraductal injection. We further show that intercalation is dependent on dynamic cytoskeletal rearrangements in both the existing monolayer and intercalating cells. These data identify luminal cell rearrangements necessary for mammary gland development and suggest a molecular mechanism for integration of cells into an existing monolayer.

## Introduction

Epithelial cells constitute the building blocks for many of the organs that comprise the animal body, and during development, epithelial tissues need to grow, remodel, and adapt to changing environments (Guillot and Lecuit, 2013; Macara et al., 2014; Pei et al., 2019). The density and stratification of epithelial cells can be modified by proliferation and extrusion, and tissue morphogenesis can be manipulated by altering cell shape. Convergent extension, which involves the exchange of cell boundaries, alters the aspect ratio of growing embryos (Sutherland et al., 2020). Intercalation of individual cells into a pre-existing epithelial layer has been described in epiboly and keel formation during zebrafish development, the developing murine ureteric bud, multiciliated cell development in *Xenopus* embryo, and in the extending branches of mammary organoids in 3D culture (Bruce and Heisenberg, 2020; Geldmacher-Voss et al., 2003; Neumann et al., 2018a; Packard et al., 2013; Stubbs et al., 2006; Wilson and Bergstralh, 2017). Moreover, re-integration of extruded cells has been observed in the *Drosophila* follicular epithelium (Bergstralh et al., 2015). However, models of epithelial cell intercalation remain rare, and the underlying mechanisms are still mostly obscure.

The murine mammary gland represents a powerful model for the investigation of various aspects of epithelial tissue morphogenesis and is particularly relevant for intercalation studies (Silberstein, 2001). The gland arises from anlagen near the nipples, and at puberty ducts begin to elongate from swollen cell clusters at their tips, called terminal end buds (TEBs). These TEBs consist of an outer layer of cap cells, which are progenitors of the myoepithelial cells that enclose the mature ducts, plus a mass of body cells, which are epithelial and give rise to luminal cells that form the inner ductal layer (Hinck and Silberstein, 2005; Sreekumar et al., 2017). Body cells are not fully polarized but form micro-lumens between cells, bordered by tight junctions, and adhere to one another through E-cadherin (Ewald et al., 2008). Somehow, the multiple layers of body cells must resolve into a single layer as the TEBs invade the surrounding fat pad of the mammary gland and the ducts behind them extend. The body cells are proliferative, but also undergo apoptosis, and a plausible model is that cell death removes excess body cells to generate the ductal lumen (Humphreys et al., 1996). Consistent with this idea, inhibition of apoptosis delays the formation of hollow ducts (Mailleux et al., 2007). However, quantitative analysis suggests that apoptotic rates are insufficient to create a single layer of luminal epithelial cells from the mass of body cells in the TEBs (Paine et al., 2016). Careful imaging studies of cultured mammary organoids showed that body cells are migratory and can intercalate into the outermost luminal layer adjacent to the basal cells, providing an alternative possible mechanism to elongate ducts (Neumann et al., 2018b).

We used published data on proliferation and apoptosis rates in TEBs and ductal elongation rates, plus experimental data on proliferation and cell division orientation in the outermost luminal layer, to model contributions to ductal elongation, and discovered that intercalation is essential and accounts for about 80% of elongation, while cell division orientation and proliferation in the outer regions of TEBs cannot account for observed elongation rates.

Intraductal injection of primary mammary epithelial cells demonstrated that intercalation into a preexisting luminal monolayer can occur in vivo. To probe mechanism, we developed a new in vitro intercalation assay using the murine Eph4 cell line and primary mammary epithelial cells. Cells in suspension when added to a confluent monolayer attach to the monolayer surface, then insert a protrusion between the monolayer cells. Spot junctions marked by ZO-1 appear around the edges of the incoming cell, which ultimately fuse with the junctions of the neighboring monolayer cells to complete the intercalation. Knockout of ZO-1 or claudin-4 in the incoming cell blocked intercalation, but surprisingly, ZO-1 knockout in the monolayer cells promoted intercalation. Actin contractility was also found to be required for efficient insertion of cells into the monolayer, but increased contractility in the monolayer inhibited intercalation. Together, these data provide a model for mammary epithelial cell intercalation that is likely applicable to many other developmental situations and may be relevant to cancer cell invasion.

## Results

### Analysis of TEBs and ductal elongation demonstrates a necessity for cell intercalation

A key question in the morphogenesis of the murine mammary gland is how the mass of body cells within the TEB self-organizes into a single layer of luminal epithelial cells to create an elongating duct. A small fraction of the body cells undergoes apoptosis, which will have a negative effect on elongation, while other cells proliferate, which will have a positive effect.

Radial intercalation of body cells into the outermost layer of luminal cells has been observed in mammary organoid formation but this process is challenging to visualize in intact mammary glands (Neumann *et al*., 2018b). While organoids are a valuable in vitro model, they do not fully reiterate the structure of the terminal end bud.

To determine if intercalation is necessary to account for the observed elongation of the mammary ductal tree, we developed a simple geometric model, based on prior work by Paine and colleagues (Paine *et al*., 2016). This earlier model examined the proliferation and apoptotic rates of both luminal and cap/basal cells within different segments of a TEB and mature duct and measured cell sizes, to determine if the observed proliferation/apoptosis rates could account for the experimentally determined rate of elongation. Our model uses the same segmentation into regions as described by Paine et al (2016), but to model the processes regulating maturation of the duct from the TEB, we focused only on the luminal cells in these regions. We segregated the TEB luminal cells into two compartments: an outermost layer that will generate the single luminal layer of the mature duct, and inner cells that form the body of the TEB (Fig. 1 A). The exterior layer of luminal cells in the TEB is continuous with the mature duct in that luminal cells in both populations contact basal cells and the extracellular matrix.

**Figure 1.**
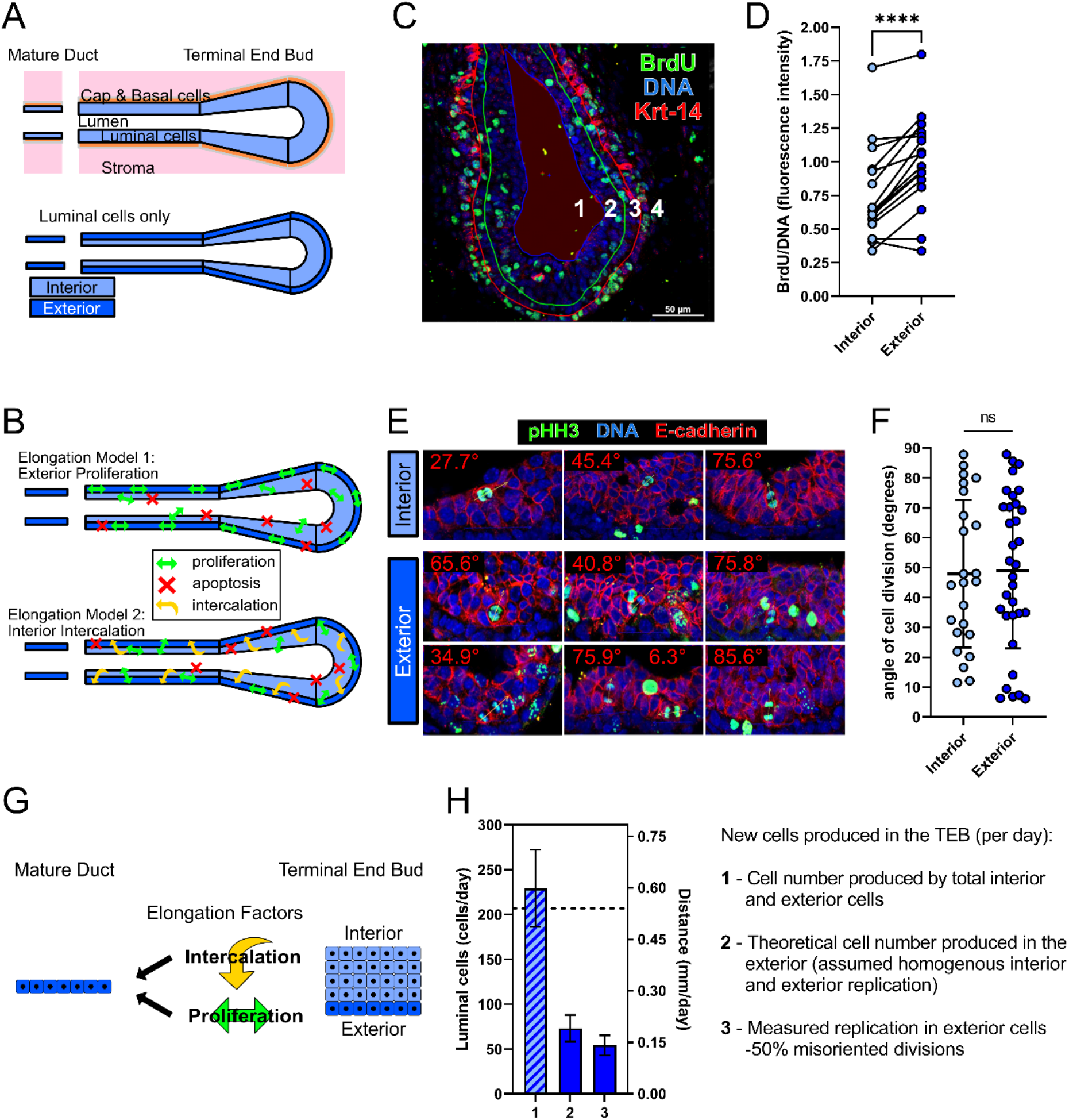
Luminal cell layer proliferation is insufficient for ductal elongation behind the TEB. A. Top: Schematic of the terminal end bud showing luminal and other cell types present, and segregated into distinct regions as described by Paine et al., 2016. Bottom: Luminal cells in the TEB with marked interior and exterior (outermost) luminal cell compartments. B. Models of luminal cell processes that might regulate elongation of the mature duct. Model 1 highlights oriented division of the exterior luminal cells contiguous with the ductal luminal layer to produce ductal extension behind the TEB. Proliferation and apoptosis in interior luminal cells in model 1 may be at steady state and interior cells contribute little to elongation. In model 2, neither cell division orientation, nor location of luminal cell proliferation and apoptosis in the TEB contribute significantly to elongation. Rearrangement by intercalation of interior body cells to the outermost layer and mature duct provides most of the elongation in this model. C. Female 6 week-old mice were administered BrdU in PBS at 100 mg/kg for 4 hrs prior to harvesting, fixing, cryo-sectioning, and staining the mammary glands. Example immunofluorescence staining of TEBs for indicated markers. Regions of interest were outlined and numbered to distinguish different regions 1: lumen, 2: interior/body luminal cells, 3: outermost luminal cells, 4: basal/cap cells. Scale bar = 50μm. D. BrdU intensity (normalized to DNA intensity) in outermost versus interior luminal TEB regions. Paired t-test, n= 16 TEBs compiled from 3 mice; p < 0.0001. E. Example TEBs stained for DNA, phosphorylated(S10)-Histone H3, and E-cadherin to highlight dividing luminal cells and determine the angle of division orientation with respect to the orientation of the outside surface of the TEB. F. Quantification of division orientations in interior and exterior TEB luminal compartments. Unpaired t-test, n= 24, 33 divisions; Mean +/- SD, p = 0.8812. G. Diagram showing how each luminal cell compartment in the TEB could contribute to the mature elongating duct. Interior cells must intercalate towards the exterior, and cells already in the outermost layer could proliferate and give rise to daughter cells in that layer. H. Quantification of new cells produced in the TEB per day (based on the geometric model with data from Paine et al., 2016 and from this study). Calculations are shown in Supp. Tables 1 and 2. Values are also represented as ductal elongation rate (mm/day) based on the average width of a mature duct cell. The dashed line at 0.54mm/day marks the average elongation of ducts per day as determined by Paine et al., 2016. Column 1 is the net total cells produced in the TEB per day in both interior and exterior luminal compartments. Columns 2 shows cells produced per day within the outermost (exterior) layer only, assuming the proliferation rate is the same in both the interior and exterior cells. The propagated uncertainty in this prediction is also shown (error bars – derivation shown at bottom of Supp. Table 1). Column 3 shows exterior layer proliferation rate calculated directly from BrdU staining in Fig. S1 B,C, which accounts for differences observed between interior and exterior replication rates (as seen in Fig. 1 C,D). Also added in this calculation is a replication correction factor of -50% for misoriented cell division (as seen in Fig. 1 E,F).

Both intercalation of cells outwardly, as well as planar-oriented division in the exterior layer, may contribute to elongation of the mammary duct. Oriented cell division is thought to be important for development of some epithelial tissues and presents a possible mechanism for elongation of the outermost luminal layer (Baena-López et al., 2005; Villegas et al., 2014). This scheme is presented as model 1 in Fig. 1B where the apoptosis of cells in the interior clears the luminal space of multilayered cells. Another hypothesis is that most of the cells contributing to elongation of the mature duct are derived from the interior subcompartment in the TEB, necessitating intercalation. In this model (model 2 – Fig. 1 B), oriented cell division in the outermost layer may contribute only a small fraction of the overall elongation.

To determine the extent to which cell division contributes to elongation, the ratio of luminal interior and exterior proliferation in the TEB were determined from sections of pubertal murine mammary glands by BrdU incorporation. Relative exterior proliferation is ∼33% greater than in the interior layers (Fig. 1 C,D, Fig. S1 A). This result might suggest that the outermost layer of luminal cells, contacting the basement membrane and continuous with the mature duct, is the primary contributor to ductal elongation. However, these cells must divide in the plane of the outermost layer to efficiently expand this layer, but stratifying, non-planar divisions have been observed previously during mammary development (Huebner et al., 2014). To determine the proportion of cells contributing to exterior elongation after division, the orientations of dividing cells in the outermost layer were measured from stained sections of mammary gland. The plane of division was found to be random (Fig.1 E,F), indicating that on average, 50% of daughter cells will be positioned in the interior after division, and in the absence of intercalation will not contribute to elongation.

To determine the extent to which TEB luminal exterior divisions contribute to elongation, we applied a geometric model, in which the dimensions of the TEB remain relatively unchanged throughout development and all excess cells generated in the TEB must contribute to the mature duct or die (Paine et al., 2016). Given these constraints, the only plausible mechanisms for elongating the duct by addition of cells are through intercalation for the interior subcompartment plus planar proliferation for the outermost layer (Fig.1 G). For each TEB region, we segregated the exterior and interior portions to first determine the extent each layer contributes to elongation, assuming the replication and death rates of cells in each compartment are the same. While the number of cells produced by the luminal cells in the entire TEB is sufficient to match the observed elongation distance (Fig.1 H – column 1), the exterior layer only provides about 30% of these cells (Fig. 1 H – column 2), suggesting most of the cells contributing to elongation must have intercalated from the interior region (Supp. Table 1).

However, these results for elongation driven by the outermost layer do not take into account division orientation or replication rate differences between the interior and exterior luminal TEB compartments observed earlier (Fig. 1 D,F). To determine how these factors alter elongation, the exterior replication rate was derived from the percent of cells with BrdU incorporation (Fig. S1 B,C, Supp.Table 2), and applied to the model. In addition, the replication rate of the outermost cells was multiplied by 0.5 to account for random division orientations. Similar to the results of the previous model (Fig. 1H – column 2), the exterior layer only provides approximately one quarter of the number of cells needed for elongation of the mammary duct when the exterior proliferation correction factors are applied (Fig. 1H – column 3). This analysis, in which only the outermost layer contributes to elongation, cannot support the extent of elongation seen during mammary gland development. Thus, our calculations identify intercalation of interior cells into the outermost layer as essential and as the main driver of mammary duct elongation.

### An in vitro assay for epithelial cell intercalation

Very little is known about the molecular mechanisms of intercalation in any system and particularly in mammalian epithelia. To address this issue, we developed a new in vitro method, initially using Eph4 murine mammary epithelial cells (Fig. 2 A). Fully confluent monolayers of mApple-labeled Eph4 cells were grown on Labtek coverglass chambers, to which unlabeled Eph4 cells in suspension were then added to simulate the multilayered state of the developing TEB (Supp. Video 1,2). The monolayers were imaged for 72 hrs, and integration was detected as the appearance of unlabeled cells within the mApple+ monolayer. The area of unlabeled cells provides a quantitative measure of successful intercalation over time (Fig. 2 B,C).

**Figure 2.**
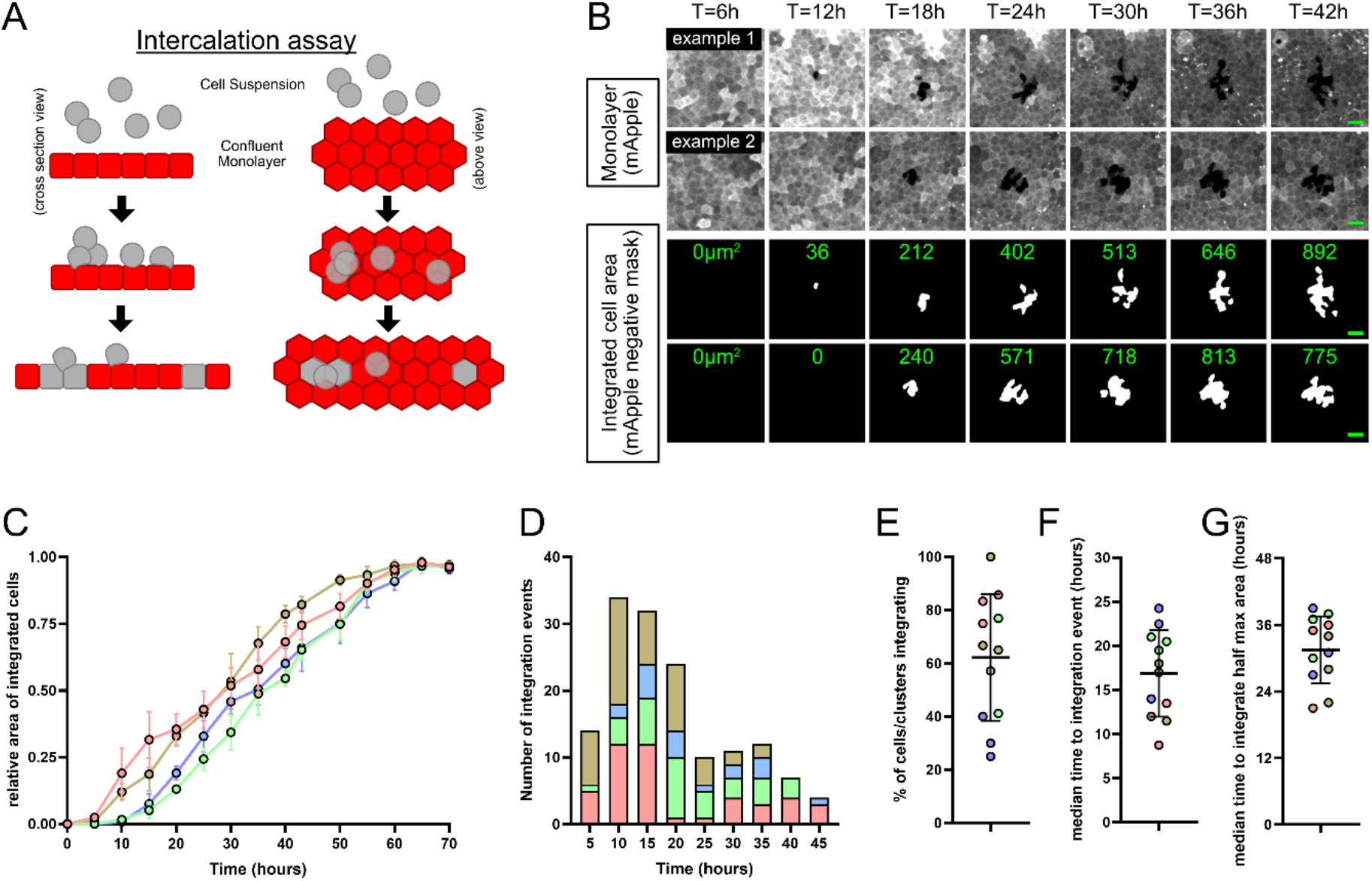
Establishment of a quantitative cell culture assay to study intercalation. A. Diagram of intercalation assay where unlabeled Eph4 cells in suspension (grey) are added to a confluent monolayer of labeled Eph4 cells (red) to simulate multilayered conditions reflective of the TEB. B. Time-lapse imaging of intercalation assay where unlabeled cells are added to a confluent monolayer of cells expressing mApple (mApple+). Area of intercalation is determined by the displacement of the mApple+ cells (and lack of fluorescence). Images are confocal projections. Lower panels show negative binary masking of the mApple+ areas, and the computed areas of integrated cells (green text). C. Measurement of the mApple-negative area over time normalized to maximum area. Values are per field of view. 4 replicates are shown. D. Number of integration events over the timelapse experiments. Each integration event is determined by the moment an mApple+ area begins to be displaced. Each color represents data from a different experimental replicate. Replicates are summed in the histogram at 5 hr increments. E. Mean percentage of cell clusters observed above the monolayer that have integrated by 70h. Determined by DIC imaging as seen in VideoS1. F. Median time until intercalation begins. Determined from Fig. 1 D. G. Median time until half of the integration area maximum is reached. Determined from Fig.1 C.

Intercalation was reproducibly detectable within 5 -10 hrs and peaked at approximately 20 hrs (Fig. 2 D,E). Of the cells or clusters observed above the monolayer, approximately 60% of these exhibited full or at least partial integration (Fig. 2 E). We noted that cells in suspension frequently intercalated as small clusters rather than as single cells and appeared to penetrate preferentially into the confluent monolayer at multi-cellular junctions. Total integration correlated positively with the number of cells in suspension added to the monolayer and correlated negatively with monolayer cell density (Fig. S2 A-D). However, if cell proliferation is inhibited using the CDK2 inhibitor Roscovitine, there is no significant effect on intercalation within 24 hrs (Fig. S2 E-H).

To determine whether primary mammary epithelial cells are capable of in vitro intercalation, we isolated cells from adult mouse mammary glands, which were grown initially as mammospheres. To select for the luminal population, cells were cultured in the absence of ROCK inhibitor, since ROCK inhibition expands the basal cell population (Prater et al., 2014). These cells were then plated on coverglass chambers to create monolayers, stained with the cell dye CFSE to label the monolayer population, and unlabeled mammary epithelial cells were then added (Fig. 3 A,B). The added cells incorporated successfully into the monolayers, with an efficiency comparable to or exceeding that of Eph4 cells at a similar monolayer density (Fig. S3 A,B). Therefore, both mammary primary cells and cell lines have the capacity for intercalation.

**Figure 3.**
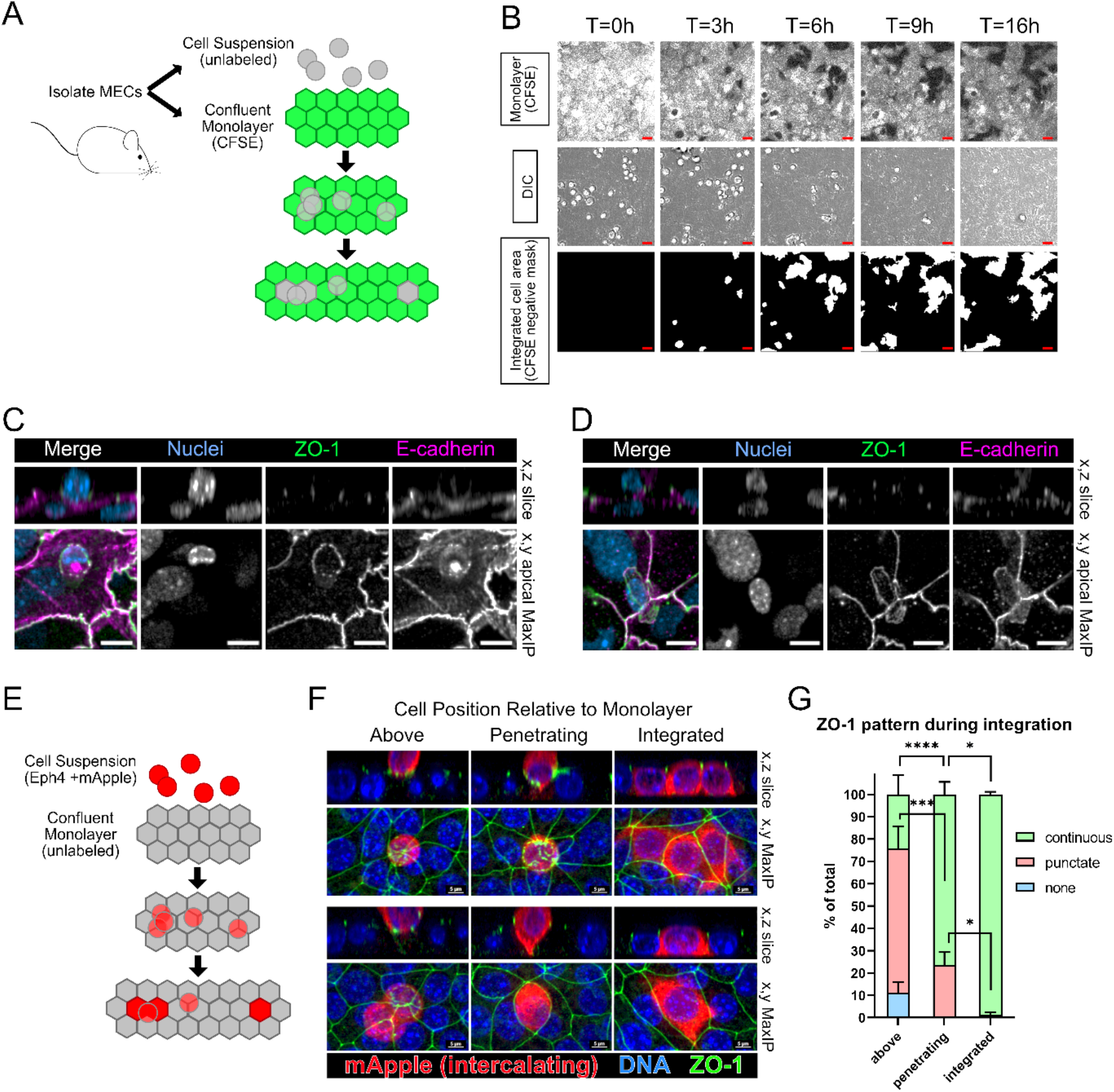
Primary mouse mammary epithelial cells intercalate through formation of junctional plaques. A. Schematic of primary mouse mammary epithelial cell (MEC) isolation, labeling and plating for intercalation assay. B. Timelapse imaging of intercalation by unlabeled MECs into confluent MEC monolayer stained with CFSE. Scale: 20μm. C - D. Immunofluorescence imaging of primary MECs undergoing intercalation, with staining for ZO-1 and E-cadherin. Cells were fixed at 24 hrs after addition of the unlabeled cells in suspension. Puncta of ZO-1 are visible at cell interface between monolayer and intercalating cells (C). D shows a cell with continuous ZO-1 staining that has moved over a multi-cellular junction. Scale: 10μm. E. Schematic of intercalation assay for F: Eph4 cell monolayer (unlabeled) and intercalating cells (mApple+). F. Eph4 cells were fixed at intervals during the intercalation assay and stained for ZO-1, then imaged as confocal z-stacks for mApple (in the intercalating cells), DNA, and ZO-1. Examples of cells displayed along the x-z and x-y axes, in 3 different states are shown: above, penetrating, or integrated into monolayer. Scale: 5μm. G. Quantification of intercalating cells in each state (above, penetrating, integrated into monolayer) with ZO-1 staining being absent, punctate, or continuous. Mean +/- SD. comparisons shown: p < 0.05.

A key step in intercalation must be the integration of the existing junctions with newly formed ones in incoming cells. To observe how these junctions assemble, intercalating cells were fixed and stained for the adherens junction marker E-cadherin and tight junction marker ZO-1. In early stages of intercalation, a discontinuous circular junction forms on the apical surface between an intercalating cell and a monolayer cell, which is positive for both ZO-1 and E-cadherin (Fig. 3 C); another example appears to be at a later stage of intercalation where the incoming cell has formed more complete junctions that have attached to existing monolayer junctions (Fig. 3 D). After complete integration, continuous junctions have formed between the intercalated and monolayer cells (Fig. S3 C).

### A role for adherens and tight junctions in epithelial cell intercalation

To further investigate the process of intercalation, we used the Eph4 in vitro intercalation system. To observe early junctional changes during this process, integration experiments were fixed 18 hrs after addition of cells to the monolayer, then immuno-stained for ZO-1 (Fig. 3 E). As with the primary cells, the Eph4 ZO-1 localization pattern is initially punctate in most incoming cells that contact the monolayer. The surface of the monolayer becomes depressed, and the ZO-1 forms a continuous junction with neighboring monolayer cells. A protrusion then penetrates between adjacent monolayer cells until it contacts the basal substrate and expands to complete the integration (Fig. 3 E,F). The maturation of the tight junction, from punctate to continuous, as intercalation occurs (Fig. 3 G), suggests that TJ dynamics could be a critical first step before cells intercalate.

To test the necessity of tight and adherens junction formation during intercalation, we generated cells null for either ZO-1 or E-cadherin using CRISPR/Cas9 gene editing (Fig. S4 A-C). The formation of intercellular adherens junctions is a universal feature of epithelial tissues, characterized by the formation of trans-interacting E-cadherin molecules on the lateral membranes. Loss of E-cadherin can result in cell extrusion from a monolayer in vitro and from mammary ducts in vivo (Shamir and Ewald, 2015; Shamir et al., 2014). We predicted, therefore, that knockout of E-cadherin in Eph4 cells added in suspension to a monolayer of WT cells would block their intercalation. To ensure a homogenous knockout, several single cell clones were isolated, and knockout was verified by immunofluorescence cell staining (Fig. S4 A-C).

We first grew cells plated sparsely and allowed them to proliferate to confluence, to determine if there were defects in monolayer formation. Loss of E-cadherin resulted in poor monolayer formation with cells lacking both tight junctions and adherens junctions (Fig. S4 A), while the absence of ZO-1 had no impact on TJ formation as determined by Claudin-4 staining, which remained junctional and at similar levels to control cells (Fig. S4 A,B). This result is consistent with previous data that ZO-1 is not essential for TJ organization (Umeda et al., 2006). E-cadherin localization was also unaffected by loss of ZO-1 (Fig. S4 A,B). There was also no significant increase in multilayering of cells in these confluent monolayers, which can be caused by extrusion or over-proliferation (Fig. S4 A,D).

When the E-cadherin-negative cells were added to WT confluent monolayers, their ability to integrate was severely impaired, demonstrating that adherens junction formation is essential for intercalation. Interestingly, the deletion of ZO-1 also impaired intercalation into a WT monolayer (Fig. 4 C-E), suggesting that it plays a key role in assembling new TJs between the incoming cell and the monolayer cells. To further probe the role of ZO-1 during intercalation, we reversed the experiment, plating ZO-1 null cells in the monolayer and adding WT cells in suspension. Remarkably, these cells intercalated more efficiently into the monolayer lacking ZO-1 than into a WT monolayer (Fig. 4 F-H). This unexpected result suggests that although ZO-1 is needed by intercalating cells to establish junctions with the monolayer, the tight junctions in ZO-1-null monolayer cells are less stable and more easily re-organized by intercalating cells.

**Figure 4.**
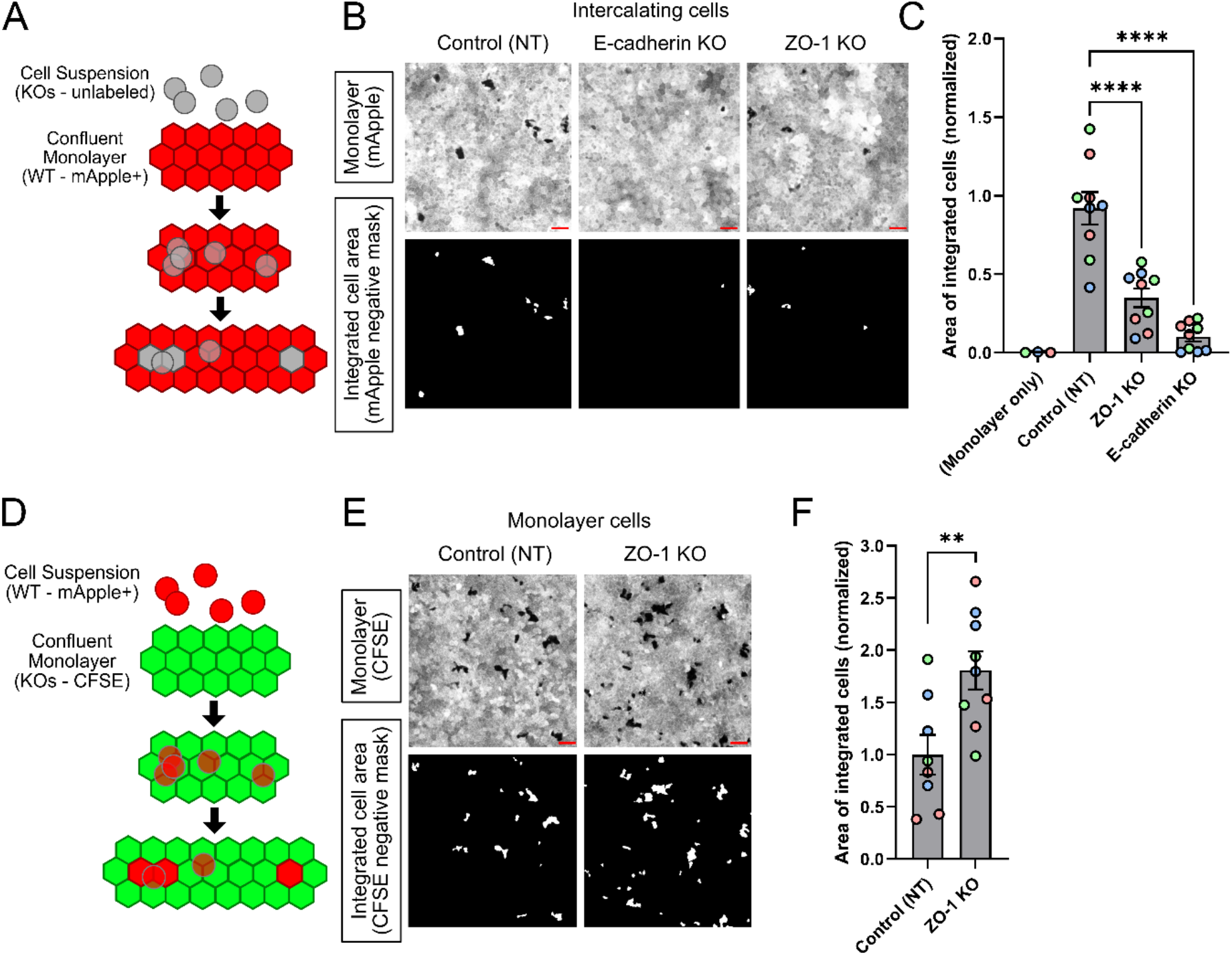
ZO-1 regulates intercalation. A. Schematic of intercalation assay for B: control monolayers and knockout cells intercalating. B. Example images of intercalation for control (NT, non-targeting gRNA), E-cadherin KO or ZO-1 KO clones. Example region shown. Scale: 50μm. C. Quantification of D: area of intercalated cells. Each clone shown in one color; n = 3 experiments. Mean +/- 1 SD; One-way ANOVA, p < 0.0001. D. Schematic of intercalation assay for E: knockout monolayers and control cells intercalating. E. Example images of intercalations for control (NT) and ZO-1 KO cells in the confluent monolayer, labeled with GFP, and WT cells added in suspension labeled with mApple. Scale bar: 50μm. F. Quantification of G showing normalized areas of intercalated cells. Each clone shown in one color. N = 3 experiments, Mean +/- 1 SD; unpaired T-test, p = 0.0080.

### Intraductally injected cells can intercalate into the luminal layer of mature ducts in vivo

We next sought to determine if intercalation into the luminal cell layer of mammary ducts can be observed in vivo, and if ZO-1 is a key regulator of this intercalation. To test this idea, a competitive transplantation assay was used where cells were introduced into adult mouse mammary glands via intraductal injection. First, mammary cells were isolated from adult mice and transduced with lentivirus to knock out ZO-1 using CRISPR (sgRNA Tjp1) and to introduce GFP as a cell marker. Other cells were transduced with lentivirus containing a control non-targeting sgRNA and expressing mCherry. Transduced cells were then cultured as mammospheres, dissociated, and combined 1:1 (GFP+: mCherry+) (Fig. 5 A). Some transduced cells were plated on coverslips and stained for ZO-1 to confirm knockout as well as to determine transduction efficiency (Fig. 5 B). The 1:1 cell mixtures were transplanted into isogenic recipient mice via injection through the nipple, and examined for GFP+ and mCherry+ cell intercalation into existing ductal monolayers. Early after injection (18 hrs), an approximately 1:1 ratio of control and ZO-1 knockout cells was detected within in the ductal lumen, confirming even numbers of each cell type were introduced (Fig .5 C,E). At 5 days post-injection, however, multiple mCherry+ control cells had integrated into the ductal luminal layer, but GFP+ ZO-1 knockout cell integration was much less apparent (Fig. 5 D,E). Intercalated cell distributions for all mice are shown in Fig. S5. This competitive transplantation assay demonstrates that intercalation can occur within mammary gland ducts in vivo, and is dependent on the major tight junction organizing protein ZO-1 to integrate cells into the mature duct.

**Figure 5.**
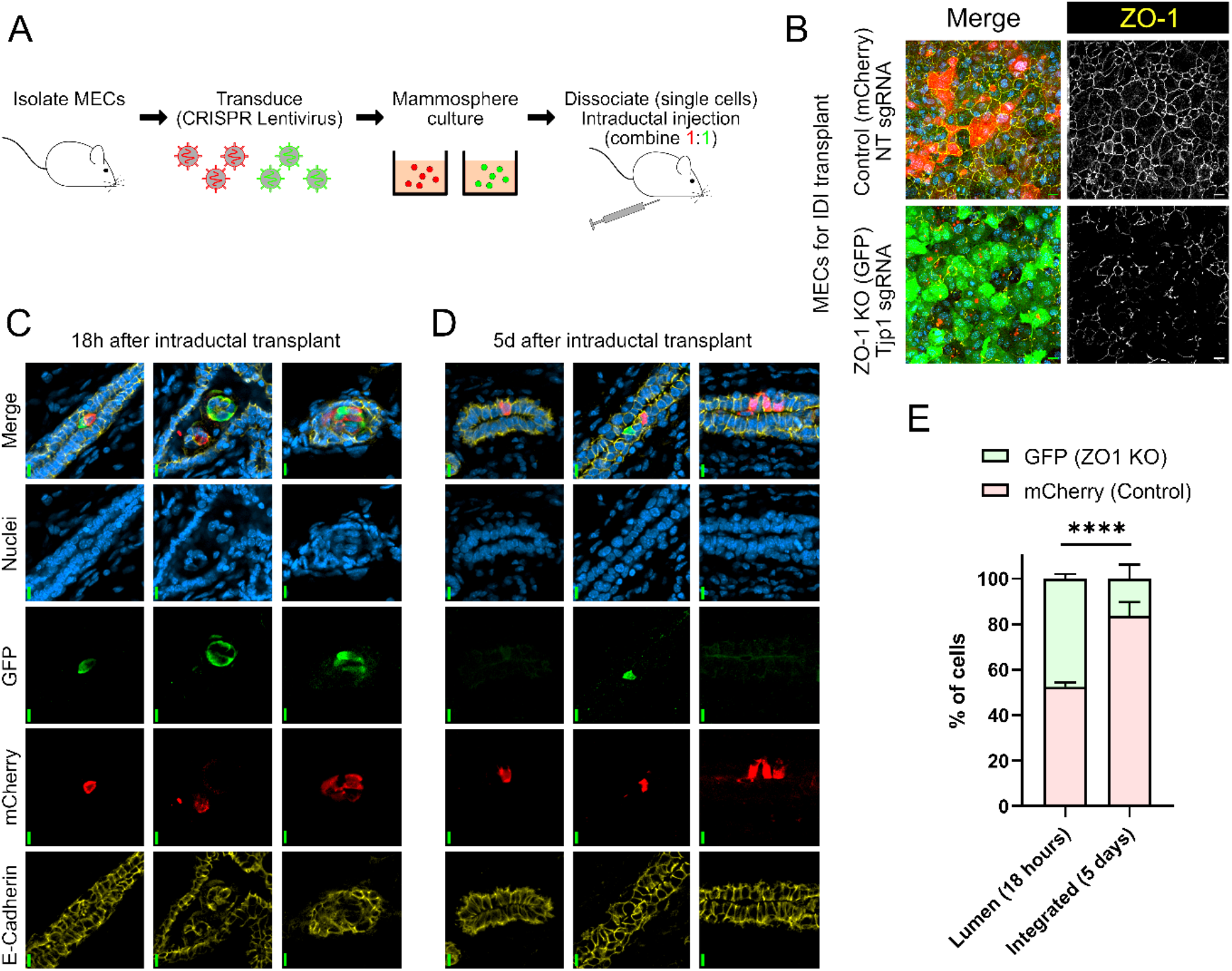
ZO-1 loss impairs intercalation of mammary epithelial cells in vivo. A. Schematic of workflow for intraductal competitive intercalation assay. Primary mammary epithelial cells (MECs) were isolated and transduced with a LentiCRISPRv2-GFP lentivirus to target ZO-1 for deletion and label cells green. Non-targeting sgRNA in LentiCRISPRv2-mCherry was used as a negative control to label WT cells red. Cells were expanded as mammospheres, then digested to single cells, mixed, and transplanted intraductally at a 1:1 ratio. B. Cells were infected with control (mCherry) or ZO-1 targeting (green) lentivirus, then were used for the intraductal injection experiments described in Fig. 6, or plated, fixed, and stained for ZO-1 to assess knockout efficiency. Scale: 10μm. C. Examples of cells present 18 h after injection in the mammary glands of recipient mice. Scale: 10μm. D. Examples of cells integrated into the existing ductal epithelium 5 days after transplantation. Scale: 10μm. E. Comparison of the distribution of cells found in the ducts at 18 hrs or 5 days post-transplantation. Per experiment: 18 hrs: 2 mice, 5 days: 5 mice. 2 experiments grouped. Fisher’s exact test, n= 413 total cells. Mean +/- SD, p < 0.0001

### Actomyosin dynamics regulate the reorganization of cells during intercalation

Actomyosin assemblies play a critical role in many cellular processes, and are likely to be involved in mammary cell intercalation as they are in other systems such as multiciliated cell development in the *Xenopus* epidermis (Murrell et al., 2015; Sedzinski et al., 2016; Ventura et al., 2022). In epithelia, the apical perijunctional actin ring supports the integrity of the connected cells in the epithelium and Myosin II also acts at this level to control contraction and rearrangement of the cells (Takeichi, 2014). To investigate the role of the actin network during intercalation, we used live imaging of cells expressing LifeAct-GFP intercalating into monolayer cells expressing mApple (Fig. 6A). Initially, an actin-rich protrusion attaches to the apical surface of a monolayer cell. This attachment expands into a circle centered between monolayer cells, as we had observed for ZO-1. The circle enlarges and eventually takes on a more polygonal shape as the cell fully integrates into the monolayer. A protrusion is often seen penetrating between the cells to reach the substratum and has begun to spread actin-rich pseudopods beneath neighboring cells (Supp. Video 3,4). We quantified the basal area occupied by incoming cells, which shows that the mean time for 50% expansion of this area is about 2 hrs (Fig. S 6A). LifeAct intensity at the apical interface between the intercalating cell and monolayer (normalized to total LifeAct intensity) remained relatively constant throughout the intercalation process even through establishment of new perijunctional F-actin rings after full integration (Fig. S 6B). At the basal side, F-actin levels increase sharply after contact with the substratum and remain at this level, while the areas occupied by integrating cells grow more slowly (Fig. S6 C).

**Figure 6.**
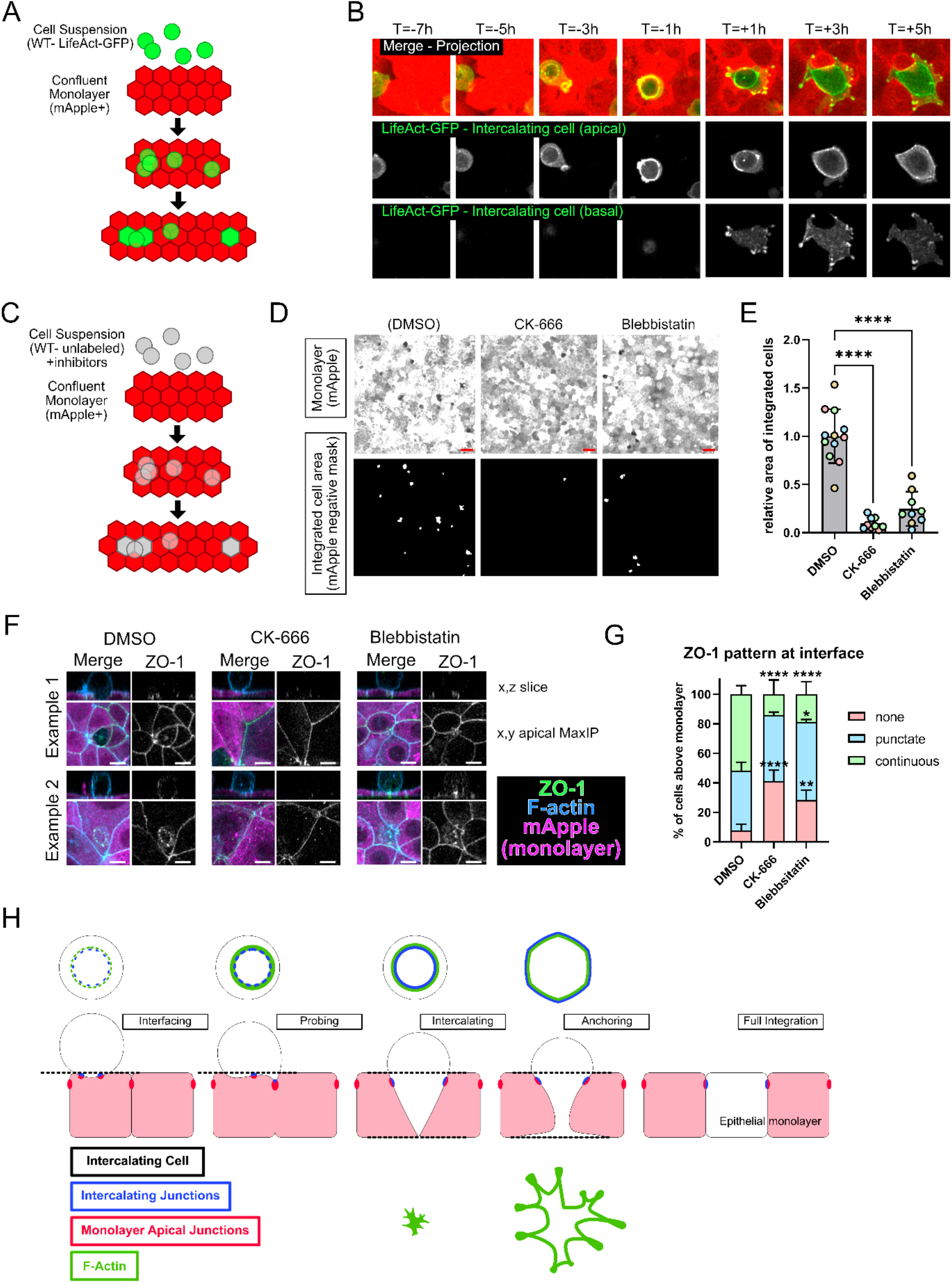
Actomyosin dynamics regulate intercalation. A. FP were added to WT monolayers expressing mApple for visualization. Live imaging with confocal microscopy. B. Stills of intercalation event with LifeAct-GFP-labeled intercalating Eph4 cell and mApple-labeled monolayer. First row: maximum intensity (MI) projection of both channels overlaid. Second row: interface of intercalating cell and monolayer, LifeAct-GFP only. Third row: basal surface, LifeAct-GFP only. Fourth row: MI projection of mApple channel only. Time normalized to initial detection of intercalation at the base of the monolayer. C. Schematic of intercalation assay for D, F: control (wildtype) mCherry+ monolayer and (wildtype unlabeled) intercalating cells added with inhibitors. D. Example images of cell intercalation and effects of CK-666 (300uM) to inhibit the ARP2/3 complex or Blebbistatin (50uM) to inhibit myosin activity. Scale bar = 50μm. E. Quantification of areas of intercalated cells. 3 experiments. One-way ANOVA, Mean +/-SD, p < 0.0001 for all comparisons shown. F. Immunofluorescence imaging of cells undergoing intercalation, with staining for ZO-1, F-actin (phalloidin), and mApple (monolayer). Cells were fixed at 16 hrs after addition of the unlabeled cells in suspension. ZO-1 pattern at the interface, excluding areas of existing monolayer junctions, was then scored. Examples are shown for each treatment with orthoganol slice and projection of the interface region. Scale: 10μm G. Quantification of intercalating cells with ZO-1 staining being absent (none), punctate, or continuous for each of the treatments shown. Mean +/- SD. comparisons shown: p < 0.05. H. Model of intercalation. Cells begin to form nascent junctions at the cell-cell interface. The actomyosin network then forms and regulates cell movement as the cell probes for an existing intercellular junction interface with which it can interact. Probing structures are seen in cell culture and similar elongated luminal cell bodies are seen in the TEB. Intercalation proceeds as the cell pushes inward and anchors itself to the substrate. Cell shape is then resolved as monolayer cells accommodate the integrated cell.

To further probe the role of the actomyosin network in this process, we tested the effects of Arp2/3 inhibitor CK-666 and Myosin II inhibitor Blebbistatin. Arp2/3 promotes branched actin formation, while the motor protein Myosin II allows for contraction along actin filaments and both of these components are involved in cell motility as well as junction assembly and rearrangement. When either Arp2/3 or Myosin II was inhibited, there was a reduction in intercalation (Fig. 6 C-E). This result shows the importance of actomyosin during intercalation; however, it is not possible to determine from these experiments if the effect of the drug is on the intercalating cells, monolayer cells, or both populations.

While these inhibitions of the actomyosin network might hamper intercalation at several stages, we first looked at the early junction rearrangement seen with ZO-1 during intercalation to see if the formation of this nascent connection was affected. Perturbations to the actomyosin network can disrupt the formation of apical junctions, and the resulting organization is often more punctate than linear (Beutel et al., 2019; Efimova and Svitkina, 2018; Engl et al., 2014). The intercalation assay was performed +/-inhibitors, and cells were fixed at 16hrs to observe early stages of intercalation. In the absence of inhibitors, ZO-1 staining at this time was mostly punctate in cells sitting atop the monolayer, with some displaying a continuous ZO-1 pattern as seen in Fig. 3 C-G (Fig. 6 F,G). When branched actin formation or myosin activity was inhibited, however, a larger fraction of cells had no ZO-1 at the interface with the monolayer. Although there was no significant difference in cells with punctate ZO-1, fewer cells were seen with continuous ZO-1 staining at the interface (Fig. 6 F,G). These observations suggest that both the initiation of puncta formation and maturation from puncta to continuous junctions are impaired, and that intercalation is hampered by the lack of continuous junction formation. We conclude that tight junction protein ZO-1 re-organization and the actomyosin network are both required for efficient intercalation (Fig. 6 H).

## Discussion

A question of fundamental importance to developmental biology is how ductal structures self-organize. The murine mammary gland provides a valuable model by which to investigate this question. The ductal tree of the gland develops over several weeks at puberty, from primordial buds attached to the nipples, and invades into the surrounding fat pad. Large terminal end buds (TEBs) at the tips of the ducts generate most of the cells that contribute to the ductal structure. A longstanding mystery, however, has been the mechanism by which the mass of body cells within the TEBs resolves into a single layer of luminal cells in the mature ducts behind the TEBs. Proposed mechanisms have implicated apoptosis of body cells to create the ductal lumen, selective proliferation and division orientation of luminal cells adjacent to the basal cell layer, and cell intercalation. However, apoptosis rates were estimated to be too small to contribute significantly to ductal organization (Paine et al 2016). Here we calculate contributions to ductal elongation of proliferation, the orientation of cell divisions within the outermost layer of body cells, and intercalation of interior cells into this outermost layer. Notably, intercalation accounts for ∼75% of the known elongation rate of the ducts, and neither division orientation nor proliferation of the outermost luminal cells contribute substantially to this rate. A consideration for the process of intercalation into the outermost luminal layer is whether migration by body cells is directional. However, if adhesions formed by the outermost cells to the extracellular matrix are more stable than the adhesions between body cells, then no directed migration is required to eventually integrate all the body cells into the outermost layer of the extending duct.

To further understand the mechanism of intercalation we developed an in vitro assay to quantify intercalation by Eph4 murine mammary epithelial cells, or by primary mammary cells. This assay also allows for live imaging and tracking of intercalating vs. monolayer populations with high resolution. We discovered that while ZO-1 is not required for normal tight junction assembly and monolayer formation in culture, it is necessary in the added cells in suspension for efficient intercalation into the monolayer. Surprisingly, however, deletion of ZO-1 from monolayer cells substantially increased rather than decreased intercalation by WT cells. We interpret this asymmetric requirement for ZO-1 as a reflection of its role in stabilizing junctions: de novo formation of spot junctions will be impaired by loss of ZO-1, but the stability of pre-existing junctions in the monolayer will likely hinder the re-organization necessary to integrate them with those of the incoming cell. High resolution imaging will be required to test this idea. Interestingly, blocking contractility or branched actin formation in all the cells, using a Myosin inhibitor or CK666, respectively, strongly inhibits intercalation.

A key question was whether our observations of intercalating Eph4 cells are applicable to actual mammary epithelia. Notably, primary murine mammary epithelial cells in culture were able to intercalate even more efficiently than the Eph4 cells; and intraductal injection of isolated mammary cells revealed in vivo intercalation into the luminal cell layer of intact ducts. Moreover, deletion of ZO-1 in the isolated mammary cells significantly inhibited in vivo intercalation, just as was seen in vitro. Intraductal injection provides, therefore, a powerful new approach to investigate the process of intercalation in an in vivo setting.

## STAR*METHODS

### KEY RESOURCES TABLE

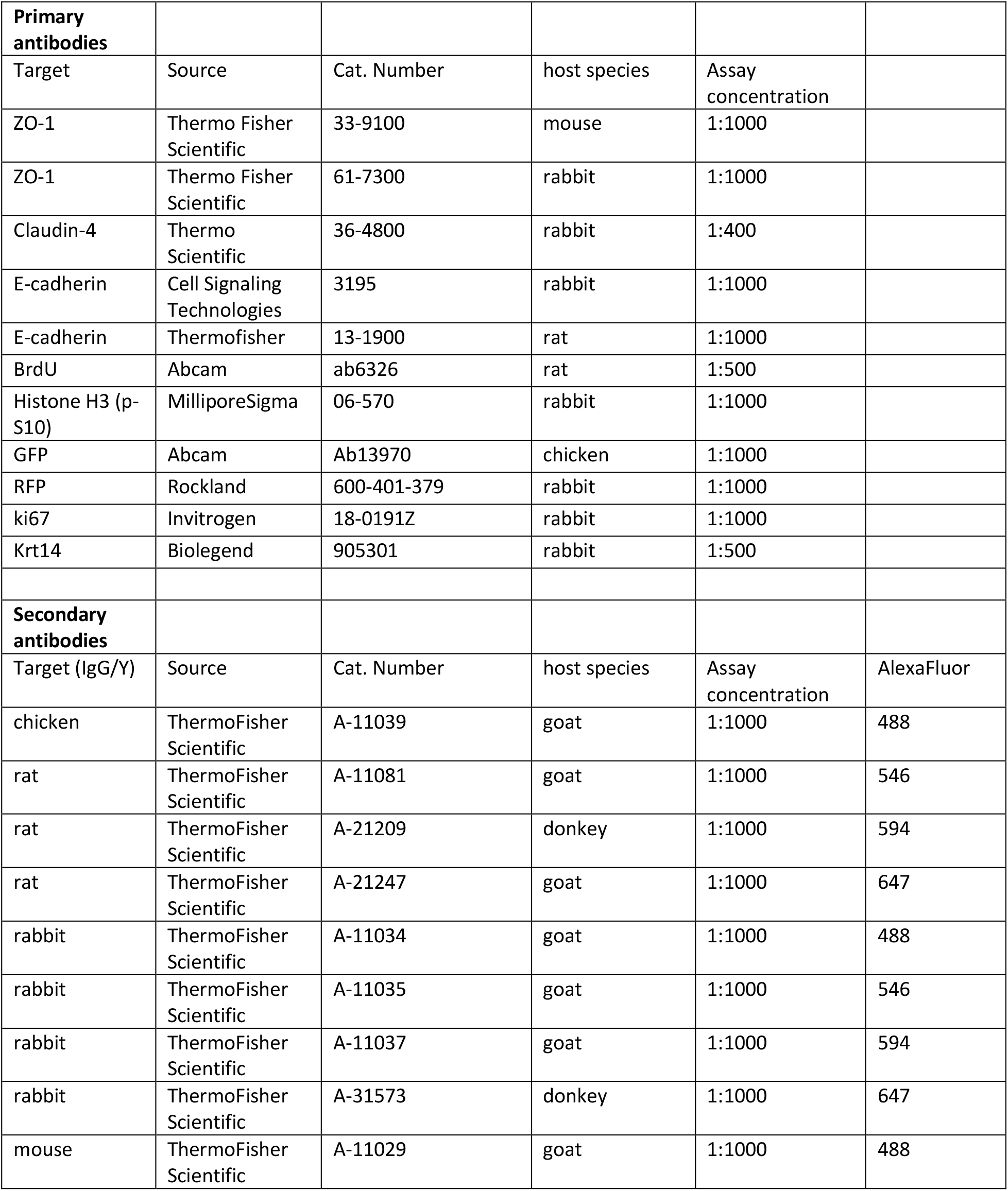
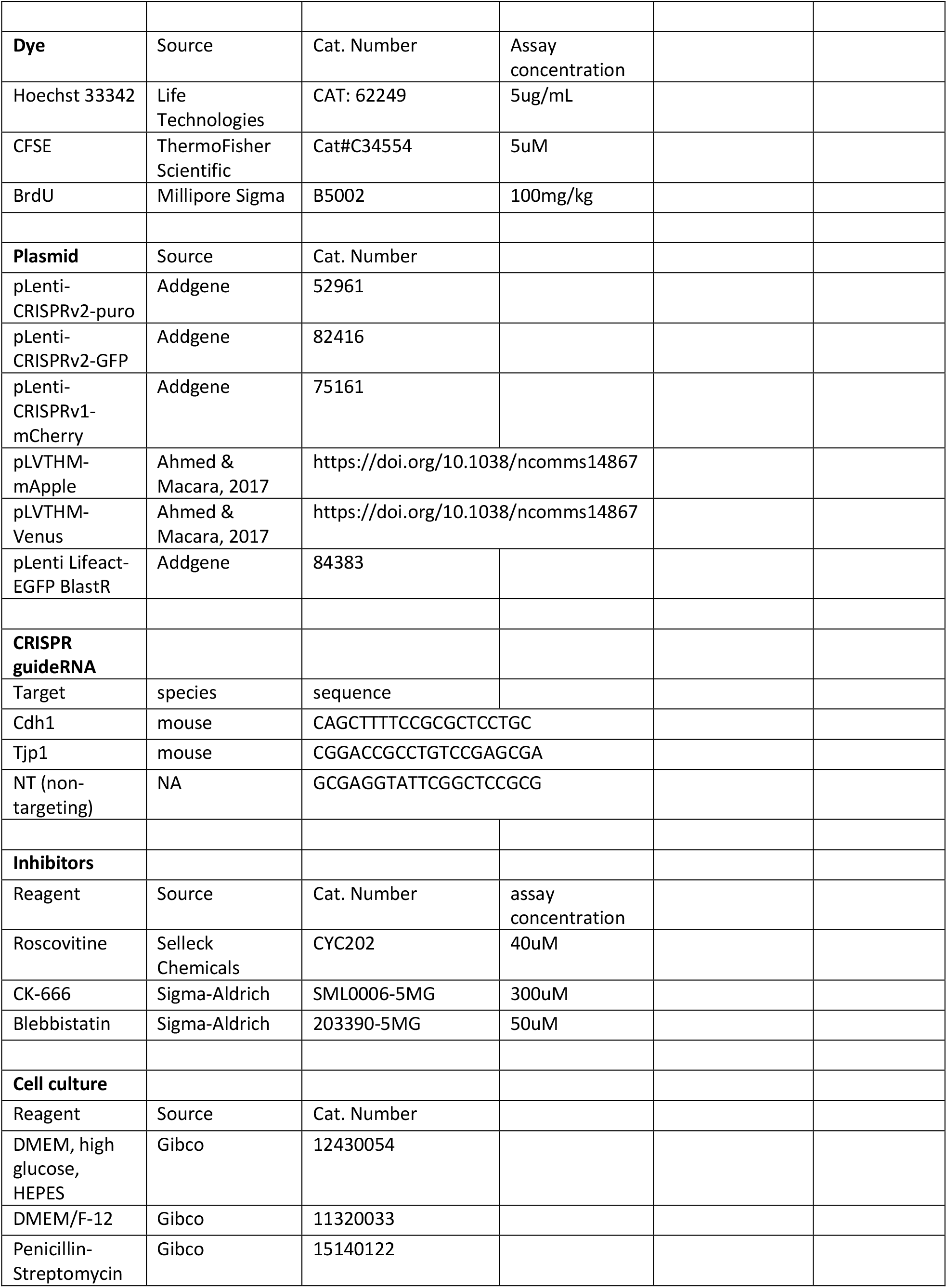
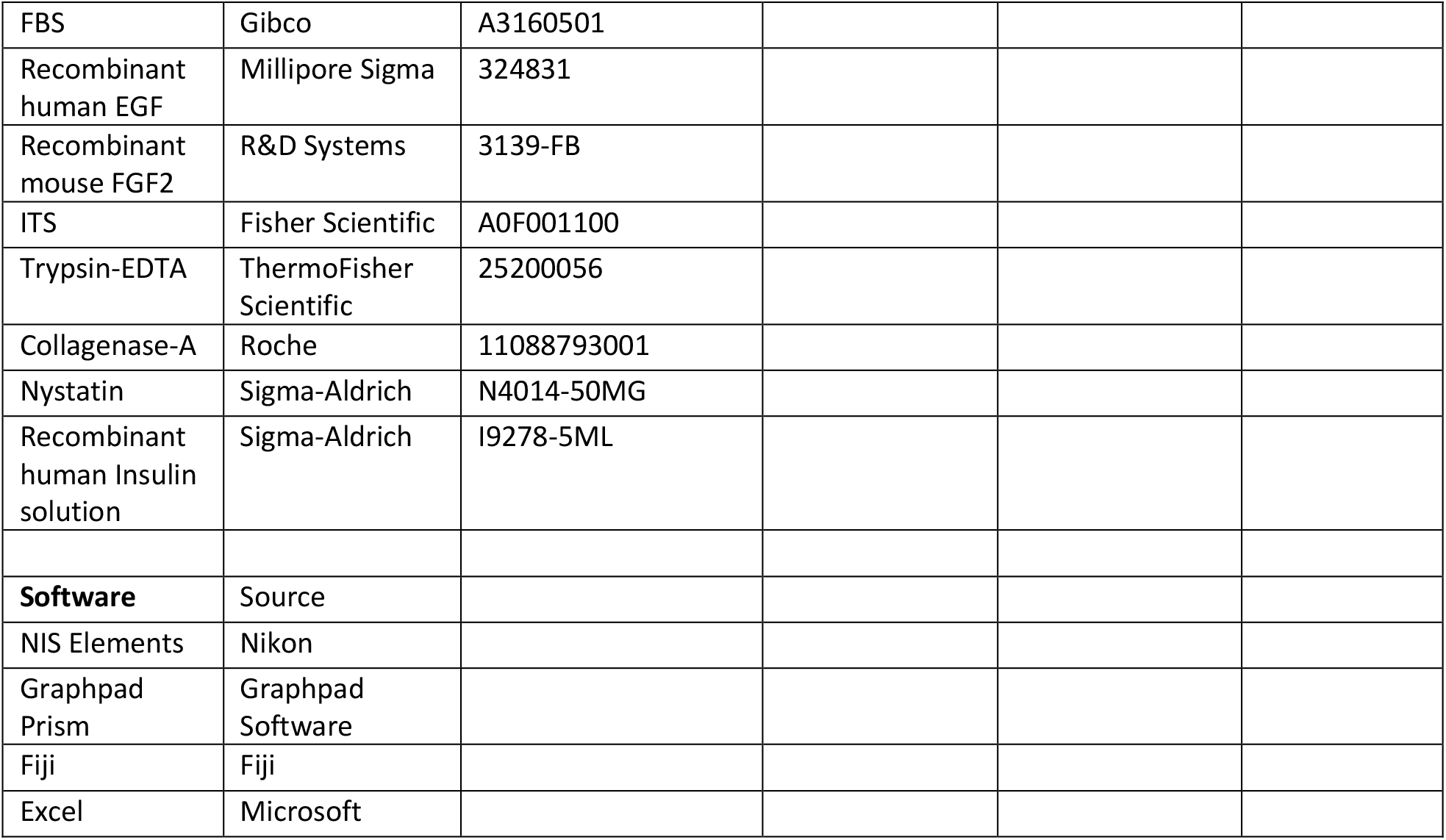

## RESOURCE AVAILABILITY

This study did not generate new unique reagents.

### Lead Contact

Further information and requests for resources and reagents should be directed to and will be fulfilled by Ian Macara

### Materials Availability

Materials such as plasmids are available upon reasonable request, or through Addgene.

## EXPERIMENTAL MODEL AND SUBJECT DETAILS

### Animals

The Vanderbilt Division of Animal Care (DAC) ensures that all mice within the Vanderbilt facility are monitored daily for health status. DAC also ensures the overall welfare of the mice, and provides daily husbandry that includes environmental enrichment, clinical care, protocol record keeping, building operations, and security. The Vanderbilt mouse facility has three experienced Animal Care Technicians who attend to the daily needs of the animals. DAC ensures that all federal, state, and university guidelines for the care and use of animals are understood and maintained. Mice were housed with a standard 12 hrs light/12 hrs dark cycle. Mice were provided normal laboratory chow and water. All mouse experiments were performed with approval from the Vanderbilt Institutional Animal Care and Use Committee.

Female FVB/NJ mice aged 6-8 weeks were used for proliferation studies related to modeling. BrdU was administered in PBS at 100mg/kg via intraperitoneal injection and mammary glands were harvested after 2 hrs. Female C3H/HeJ mice aged 8 wks or older were used as donor and recipient for intraductal transplantation experiments. All mice were acquired from The Jackson Laboratory.

### Isolation and culture of primary cells

The 4^th^ pair mammary glands were isolated from adult female mice, minced with scissors, and digested in DMEM/F12, 2mg/mL collagenase, 100U/mL Penicillin and Streptomyocin, 600U/mL Nystatin, and 5μg/mL insulin for 1 hr shaking at 37°C. The resulting pellet was then treated with 2U/mL DNase for 5 min, washed 5x with DMEM/F12, then digested to single cells with 0.25% Trypsin for 12 min shaking at 37°C. Single cells were then strained through 40μm pore strainer. Single cells were transduced and grown in low-attachment plates in DMEM/F12, 5ng/mL EGF, and 1x ITS. Aliquots of cells were plated onto cover-glasses and allowed to adhere and form monolayers before fixing and immunostaining to determine the percentage of cells in each condition that were transduced and the efficiency of the CRISPR knockout. The mammospheres (low-attachment culture) were digested using Trypsin at 37°C to obtain single cells before being combined 1:1 (control:knockout) to be transplanted into the mammary duct. For primary cell in vitro intercalation assays, 150×10^3 single cells were plated onto 8-well chambered coverglasses and allowed to proliferate to confluence for 1-3 days. Monolayer cells were then stained for 10 min at 37°C with CFSE (2.5ug/mL in DMEM/F12), washed 3 times for 5 min each with DMEM/F12 plus 5% FBS. Unlabeled primary cells were then added and live imaging was started within 30 minutes to observe intercalation.

### Cell lines

Mouse mammary EpH4 cells were provided by Dr. Jürgen Knoblich (Institute of Molecular Biotechnology, Vienna, Austria). EpH4 cells were cultured in Dulbecco’s Modified Eagle Medium (DMEM) (ThermoFisher Scientific, Waltham, MA) supplemented with 10% Fetal Bovine Serum (FBS) (R&D Systems, Minneapolis, MN), and incubated at 37°C and 5% CO2.

## METHOD DETAILS

### TEB luminal elongation model and calculations

A geometric two-dimensional model of murine mammary TEBs was based on that described by Paine et al (2016), in which the TEB was subdivided into several regions. For the purposes of our analysis, however, we ignored the basal layer of cap cells, and segregated the luminal body cells into two compartments, an outermost (exterior) layer of luminal cells that is adjacent to the basal cap cells, and an interior compartment that comprises all other body cells in the TEB. The area of each luminal compartment was then calculated from measured parameters (Paine et al 2016).

Area of the exterior layer, A_E_ = TEB perimeter x cell height Interior area A_I_ = total TEB area, A_T_ – A_E_

We then determined the ratios A_I_/A_T_ and A_E_/ A_T_. These ratios were multiplied by the total luminal cells per region and summed to calculate the numbers of exterior and interior cells:

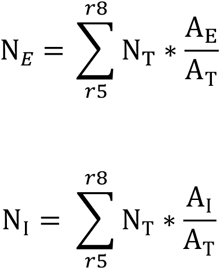

To determine the contribution of just the exterior cells to ductal elongation, their rates of proliferation and death rates were calculated (Table 1). Corrections to the proliferation rate, r, were determined as shown in Fig 1. Correction 1 was for the higher observed proliferation rate, r_E_, of the exterior cells relative to interior cells. (Corrected r_E_ = 1.25* r_E)_. Correction 2 was for the random orientation of cell division in the outermost layer. (Corrected r_E_ = 0.5*r_E_). Both corrections were applied in the final calculations.

#### Determining the proliferation rate of the outermost luminal layer in the TEB

We empirically determined the rate of division in the exterior luminal TEB by BrdU incorporation as compared to that of interior luminal cells. BrdU was administered to 6wk-old mice and the percent of BrdU+ luminal cells in the outermost layer and inner cells was determined. The replication rate was then extrapolated from this data as described previously in Paine et al., 2016, based on the duration of S-phase, when cells are receptive to BrdU labeling, being approximately 6 hrs.

### Plasmid construction and lentivirus production

The sgRNAs used in this study are listed in the Resources Table. gRNAs were designed using CHOPCHOP. sgRNAs were cloned into lentiCRISPR v1 or v2 vector at the BsmBI restriction site using Zhang lab protocol (Shalem et al., 2014; Sanjana et al., 2014). pLVTHM-mApple was generated as described in Ahmed et al., 2017. pLVTHM-Venus was generated as described in McCaffrey et al., 2009.

293T cells were cultured in DMEM, 10% FBS, and 100U/mL Penicillin and Streptomyocin. To produce lentivirus, 293T cells were transfected with packaging plasmids psPax2 and pMDG2-VSVG along with the desired plasmid to be packaged in the lentiviral genome. This was done using calcium phosphate transfection. Medium was changed after 18 hrs, and virus-containing media was collected after 36 hrs. In some cases, virus-medium was concentrated using Amicon 100k centrifugal filters.

### In vitro intercalation assay

Eph4 cells were cultured in DMEM, 10% FBS (fetal bovine serum), and 100U/mL Penicillin & Streptomyocin. For intercalation assay, 150×10^3 single cells were plated onto 8-well chambered coverglasses. 24 hrs later, other cells were then dissociated to single cells with Trypsin and washed before being added to the previously plated monolayers. Integration of the intercalated cells into the monolayer was then determined by imaging the area displaced in the monolayer after 24 hrs. In all assays where area was determined, monolayer cells expressed a fluorescent marker. Thresholding was used to determine the area where monolayer signal was displaced, and the total area of integration was summed. Figures include example images of the monolayer fluorescence, masks of the determined area of integration, and normalized area of integration quantifications. For many experiments, co-stain with either Hoechst or differentially labeled cell populations were used to ensure areas where monolayer cells were displaced were indeed intercalated cells.

### In vivo intercalation assay

Single mammary epithelial cells were transplanted into mammary ducts via the nipple of anesthetized mice. Cells either expressed: CRISPR-mCherry-non-targeting-sgRNA or CRISPR-GFP-Tjp1-sgRNA. Labeled cells were combined in a 1:1 ratio and 5-10 x10^3 labeled cells in 10μL of DMEM/F12 (basal media – no growth factors) were injected with a custom Hamilton syringe (blunt end style: 3, gauge: 30, needle length: 15mm). Transduction efficiency was less than 100% and as a result approximately 5-10 x10^3 unlabeled wildtype cells were also injected. Mice were harvested 18 hrs or 5 days after transplantation and the composition of cells present in the lumen or integrated in the duct, respectively, was quantified.

### Immunostaining

Cells were fixed with 4% paraformaldehyde for 10 min, or overnight for tissue. Tissue was then placed in 30% sucrose solution for at least 24 hrs until embedding and cryo-sectioning. Samples (cells and tissue) were blocked and permeabilized with 5% normal goat serum in PBS with 0.2% TritonX-100. Primary or secondary antibodies were incubated for 1 hr with cells or overnight at 4°C for tissue sections. Hoechst or phalloidin stains were added during secondary antibody incubation. Slides were mounted with Fluoromount-G.

### Confocal microscopy and image processing

All imaging was performed using a Nikon A1R scanning confocal microscope and analysis was done with Nikon Elements software. Live cell imaging was performed in a TOCRIS chamber at 37 °C with 5% CO2. Imaging was performed at intervals as specified per experiment. Generally, intervals for long term intercalation experiments were 30min, and were 15-20min for shorter intercalation experiments. Confocal images were collected with z-steps ranging from 1.5μm for intercalation assay field analysis or 0.0 - 1μm step for higher magnification analysis of cell junction staining. Objectives used: 20x, 40x, 60x. Maximum intensity projections (IP) were created for display of intercalation assay field analysis and for other staining experiments. TEB images are individual sections and not maximum intensity projections.

### Statistical analysis

Statistical tests performed are described in the figure legends for each graph with comparisons. In general, unpaired t-test, paired t-test, one-way ANOVA, and Fisher’s exact test were performed in GraphPad Prism. Graphs shown also indicate error bar definitions in figure legends (SD, SEM, etc.). p values are listed in the figure legends for all comparisons shown on each graph.

**Figure S1.**
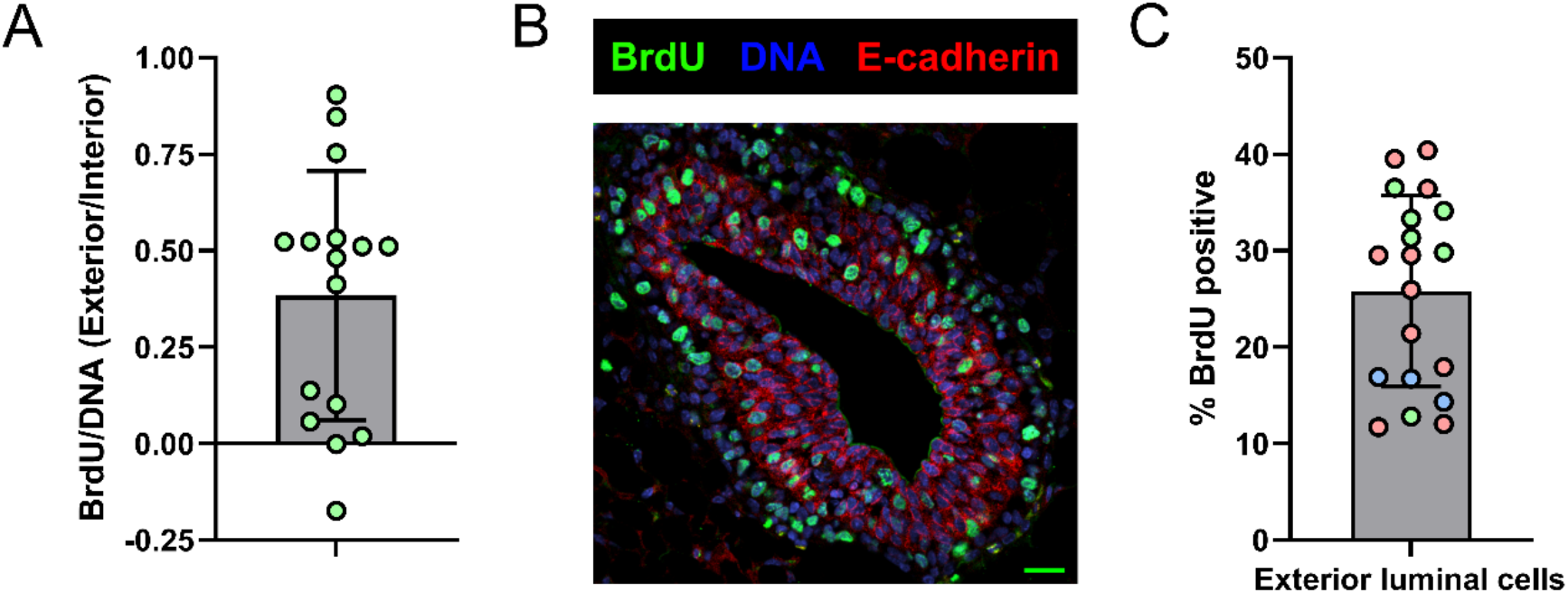
Models of the TEB and mechanisms contributing to elongation. A. Ratio of relative BrdU/DNA intensities in exterior vs interior TEB luminal compartments. Mean +/- SD B. Female 6 week-old mice were administered BrdU in PBS at 100 mg/kg for 2 h, prior to harvesting, fixing, cryo-sectioning, and staining the mammary glands. Example immunofluorescence staining of TEBs for indicated markers. Scale bar = 20μm. C. BrdU incorporation in the exterior luminal cell regions only (determined from Fig. S1 B) Measured as percent of nuclei with positive BrdU staining. 3 mice, 19 TEBs. Mean +/- SD.

**Supplementary Table 1.**
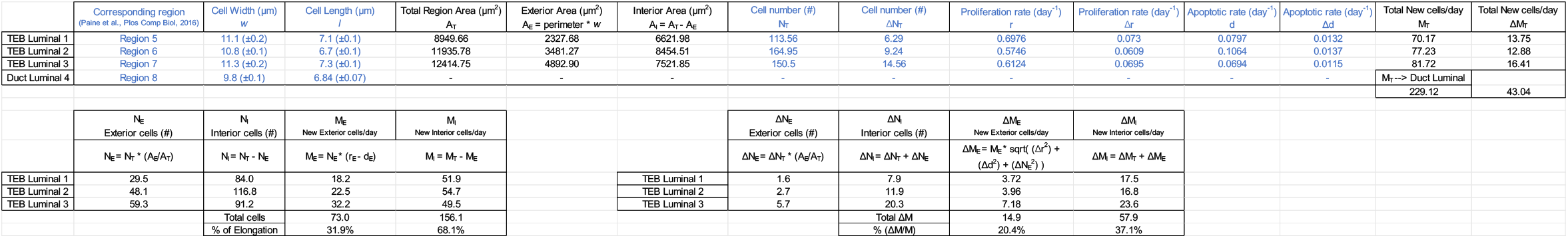
Values and calculations used to determine TEB exterior and interior luminal cell contribution to mature duct. Values measured by Paine et al. (2016) are highlighted in blue. Derivation of the uncertainty in this model is also shown in bottom right table.

**Supplementary Table 2.** Data showing %BrdU positive exterior cells converted into replication rate for outermost (exterior) luminal cells. These values along with the correction factor for random division orientation were then used to determine the number of cells generated in the exterior layer per day, and the ductal elongation rate.

| Mouse | TEB | BrdU positive (%) | Proliferation rate (day<sup>-1</sup>)<br>r<sub>E</sub> | Mature Duct Cell Length<br>(μm) l | Luminal cells (N<sub>E</sub>) | Apoptotic rate (day<sup>-1</sup>)<br>(Luminal Average) d<sub>E</sub> | M<sub>E</sub><br>New Exterior cells/day |  |  |  |
| --- | --- | --- | --- | --- | --- | --- | --- | --- | --- | --- |
|  |  |  |  |  |  |  | M<sub>E</sub> = N<sub>E</sub> * ((r<sub>E</sub> * 0.5) - d<sub>E</sub>) | average M<sub>E</sub><br>(mouse) | Duct Elongation Length<br>(mm) L = M<sub>E</sub> * (l / 2) | % of total elongation |
| 1 | 1 | 12.0% | 0.480 | 6.840 | 137.0 | 0.085 | 21.2 | 60.7 | 0.0725 | 9.3% |
| 1 | 2 | 17.9% | 0.718 | 6.840 | 137.0 | 0.085 | 37.5 |  | 0.1283 | 16.4% |
| 1 | 3 | 39.5% | 1.581 | 6.840 | 137.0 | 0.085 | 96.6 |  | 0.3305 | 42.2% |
| 1 | 4 | 11.7% | 0.467 | 6.840 | 137.0 | 0.085 | 20.3 |  | 0.0694 | 8.9% |
| 1 | 5 | 21.4% | 0.854 | 6.840 | 137.0 | 0.085 | 46.9 |  | 0.1602 | 20.4% |
| 1 | 6 | 36.4% | 1.455 | 6.840 | 137.0 | 0.085 | 88.0 |  | 0.3008 | 38.4% |
| 1 | 7 | 40.4% | 1.615 | 6.840 | 137.0 | 0.085 | 99.0 |  | 0.3385 | 43.2% |
| 1 | 8 | 25.9% | 1.035 | 6.840 | 137.0 | 0.085 | 59.2 |  | 0.2026 | 25.9% |
| 1 | 9 | 29.5% | 1.182 | 6.840 | 137.0 | 0.085 | 69.3 |  | 0.2369 | 30.2% |
| 1 | 10 | 29.5% | 1.180 | 6.840 | 137.0 | 0.085 | 69.2 |  | 0.2366 | 30.2% |
| 2 | 12 | 33.3% | 1.333 | 6.840 | 137.0 | 0.085 | 79.7 | 69.5 | 0.2724 | 34.8% |
| 2 | 13 | 29.8% | 1.191 | 6.840 | 137.0 | 0.085 | 69.9 |  | 0.2392 | 30.5% |
| 2 | 14 | 34.1% | 1.366 | 6.840 | 137.0 | 0.085 | 81.9 |  | 0.2800 | 35.7% |
| 2 | 15 | 36.5% | 1.459 | 6.840 | 137.0 | 0.085 | 88.3 |  | 0.3020 | 38.5% |
| 2 | 16 | 12.8% | 0.512 | 6.840 | 137.0 | 0.085 | 23.4 |  | 0.0799 | 10.2% |
| 2 | 17 | 31.3% | 1.250 | 6.840 | 137.0 | 0.085 | 73.9 |  | 0.2529 | 32.3% |
| 3 | 19 | 14.3% | 0.571 | 6.840 | 137.0 | 0.085 | 27.5 | 32.1 | 0.0940 | 12.0% |
| 3 | 20 | 16.9% | 0.677 | 6.840 | 137.0 | 0.085 | 34.7 |  | 0.1187 | 15.1% |
| 3 | 21 | 16.7% | 0.667 | 6.840 | 137.0 | 0.085 | 34.0 |  | 0.1163 | 14.8% |
|  |  |  |  |  |  | Average | 59.0 | 54.1 | 0.2017 | 25.7% |
|  |  |  |  |  |  | SD | 27.11054251 |  | 0.092718055 | 11.8% |
|  |  |  |  |  |  | SEM |  | 11.3095674 |  |  |

**Figure S2.**
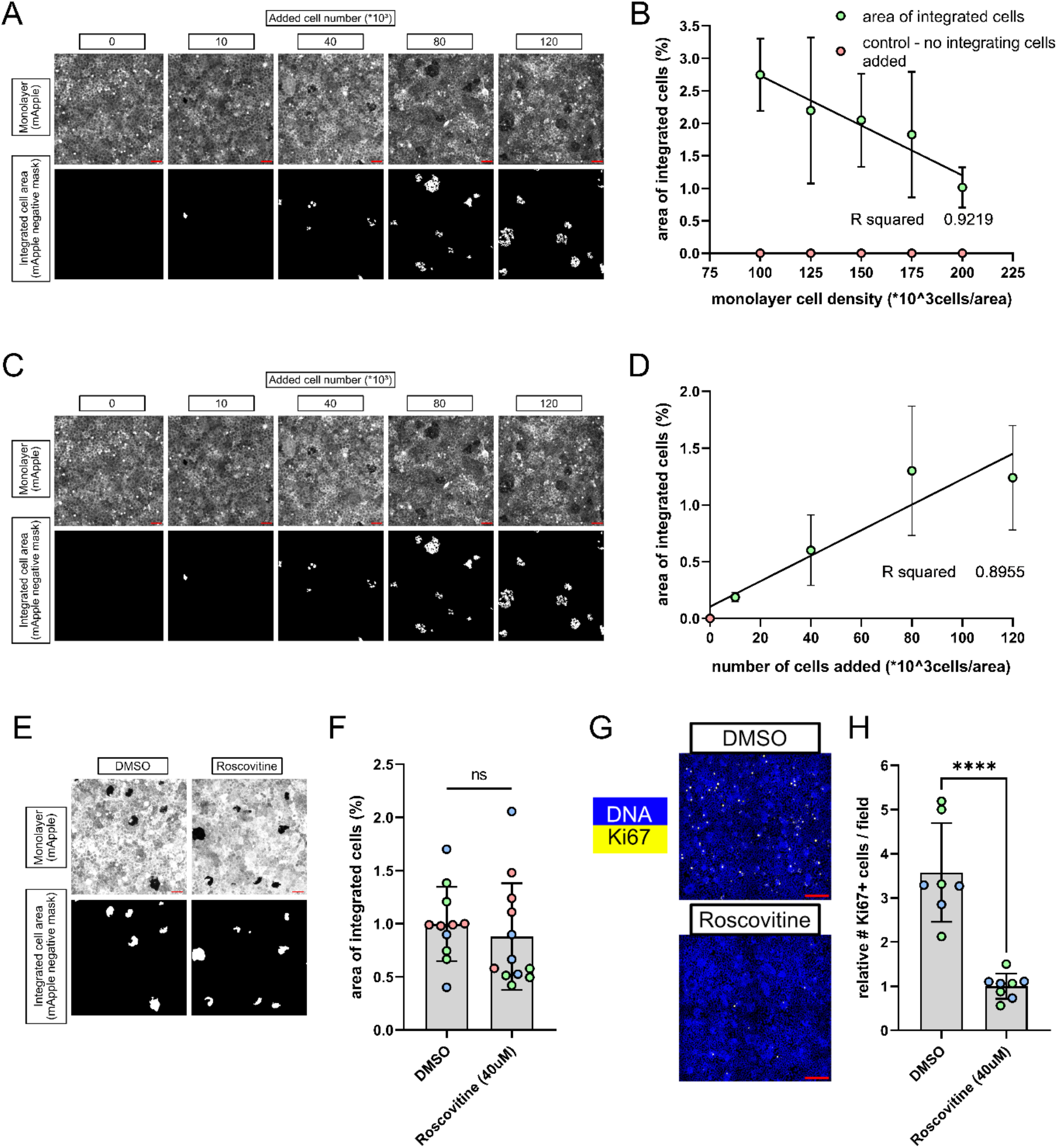
Cell density but not cell division alters intercalation dynamics. A. Intercalation assay with variable monolayer cell densities at plating. Scale: 50μm. B. Plot of integrated cell area (as percent of total monolayer area) versus monolayer density. 1 example experiment shown. Mean +/- SD. C. Intercalation assay with variable added (intercalating) cell number and fixed monolayer density. Scale: 50μm D. Plot of integrated cell area (as percent of total monolayer area) versus added cell number. 1 example experiment shown. Mean +/- SD. E. Intercalation assay performed after treatment of cells with 40 μM Roscovitine, to inhibit cell division, or vehicle (DMSO). Example field of mApple+ monolayers (top) with areas displaced by unlabeled intercalating cells shown below as inverse binary masks. Scale: 50μm. F. Areas of integration from Fig S2 E. Unpaired t-test. Mean +/- SD, n = 3 experiments, p = 0.5303. G. Field of Eph4 cells from intercalation assay in Fig. S2 E stained for DNA and Ki67 as a proliferation marker. Scale: 200μm. H. Normalized number of Ki67-positive cells per field. 2 experiments. Unpaired t-test. Mean +/- SD; p < 0.0001.

**Figure S3.**
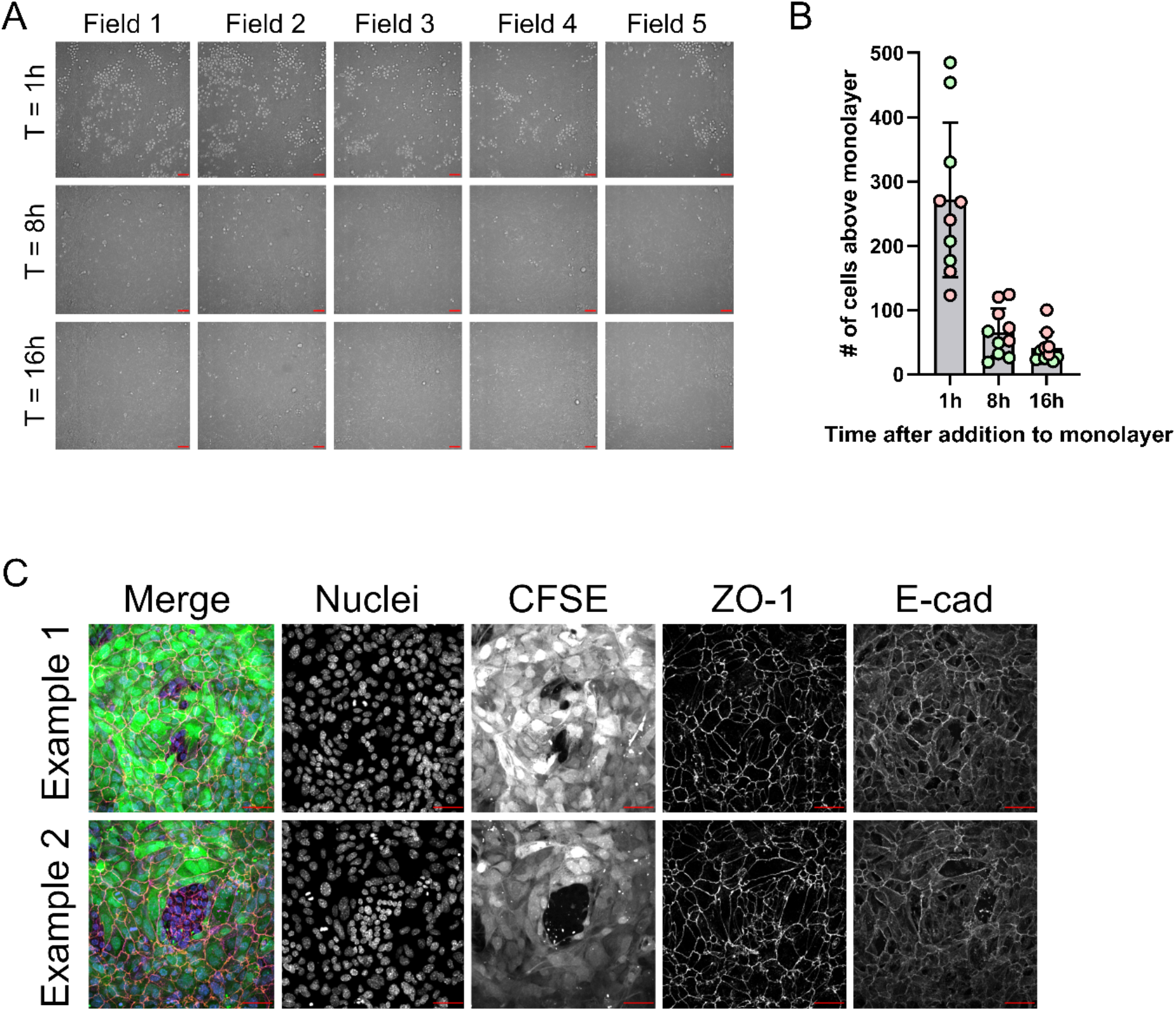
Primary cell intercalation quantification and. A. Timelapse imaging of DIC channel only. Cell clusters sitting atop monolayer are apparent. 5 example fields. Scale: 50μm. B. Total number of cell clusters located above the monolayer. 2 experiments graphed. C. Intercalation assay fixed after 24hrs and stained for junctional markers, showing full integration of intercalating cells into the monolayer. Scale: 20μm.

**Figure S4.**
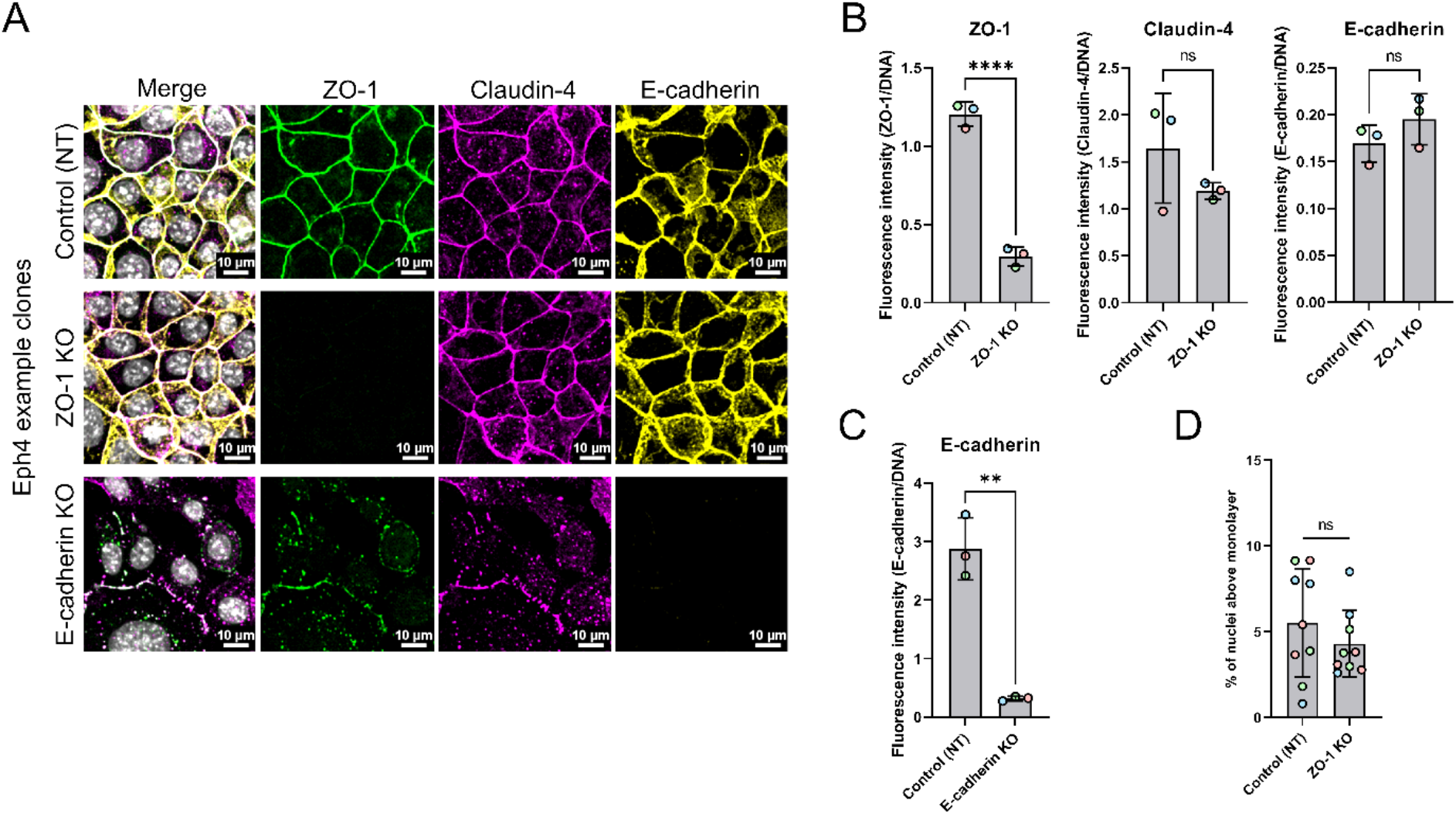
ZO-1 loss does not impair monolayer organization and does not affect junctional recruitment of Claudin-4. A. The same number of cells were plated for each clone and fixed at confluent density. Cells were then immunostained for ZO-1, Claudin-4, E-cadherin, and DNA. Deletion of ZO-1 does not alter the location of either Claudin-4 or E-cadherin to intercellular junctions, but deletion of E-cadherin disrupts tight junctions. Immunofluorescence of knockout clones after expansion to confluent monolayer. One example clone for each knockout (ZO-1 and E-cadherin) is shown. Scale: 10μm. B. Quantification of fluorescence intensities for total ZO-1, Claudin-4, and E-cadherin in fields of control (non-targeting sgRNA) and ZO-1 knockout (Tjp1 sgRNA) clones. Intensity for each channel was determined for a random field for each clone. Intensities for each marker were normalized to DNA intensity per the same field to account for subtle cell number differences. Mean +/-SD. unpaired T-test. p < 0.0001, p = 0.2514, p = 0.2519 C. Comparison of E-cadherin staining of control (non-targeting sgRNA) clones and E-cadherin knockout (Cdh1 sgRNA) clones. Mean +/-SD. unpaired t-test. p = 0.0011 D. Multilayering analysis of control and ZO-1 knockout clones as measured by percent of nuclei above the monolayer compared to total nuclei. Mean +/-SD. unpaired t-test. p = 0.3384

**Figure S5.**
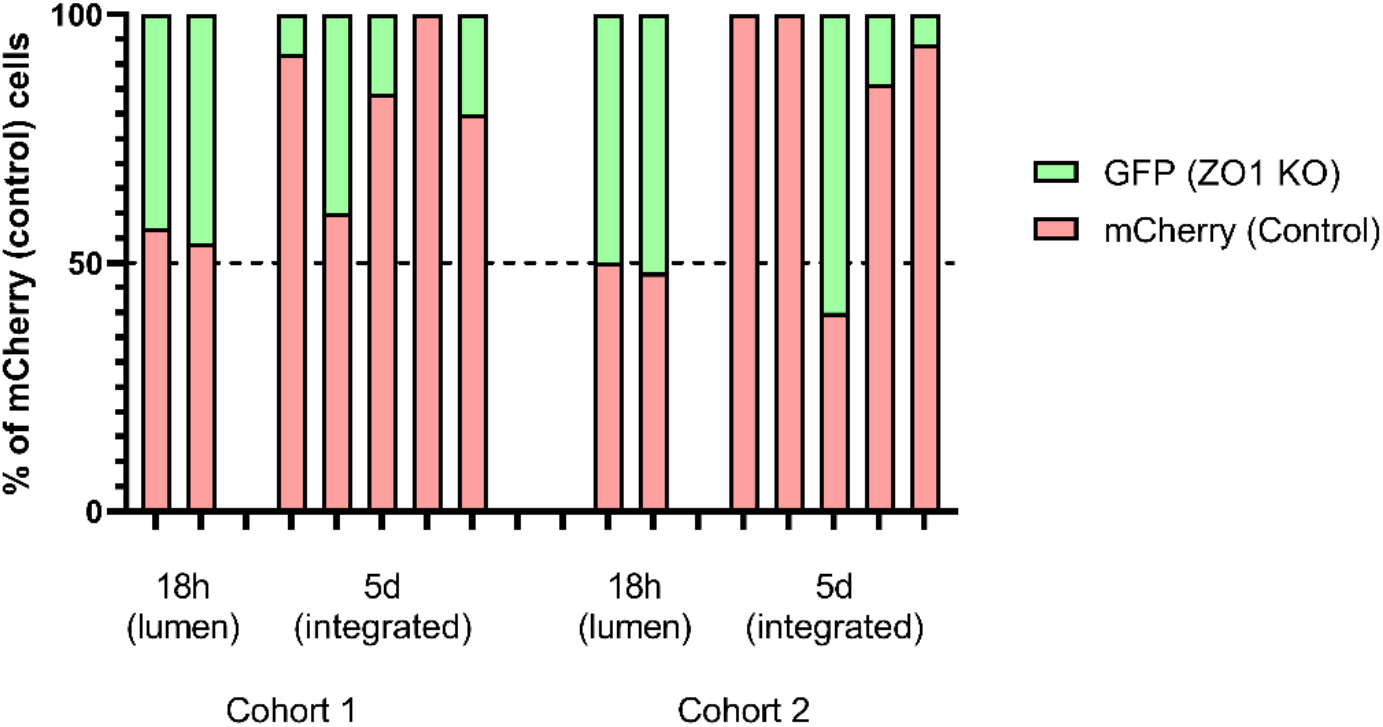
ZO-1 knockout verification in primary mammary epithelial cells, individual mice from transplants. A. Distribution of cells found in each mouse after intraductal injection. % mCherry control cells shown for simplicity (of total mCherry and GFP cells). The first 2 columns of each cohort are mice taken soon after injections to verify 1:1 distribution in the lumen (50% mCherry control cells). Next columns are mice taken after 5 days to allow cells in lumen to intercalate. Dotted line drawn at 50%.

**Figure S6.**
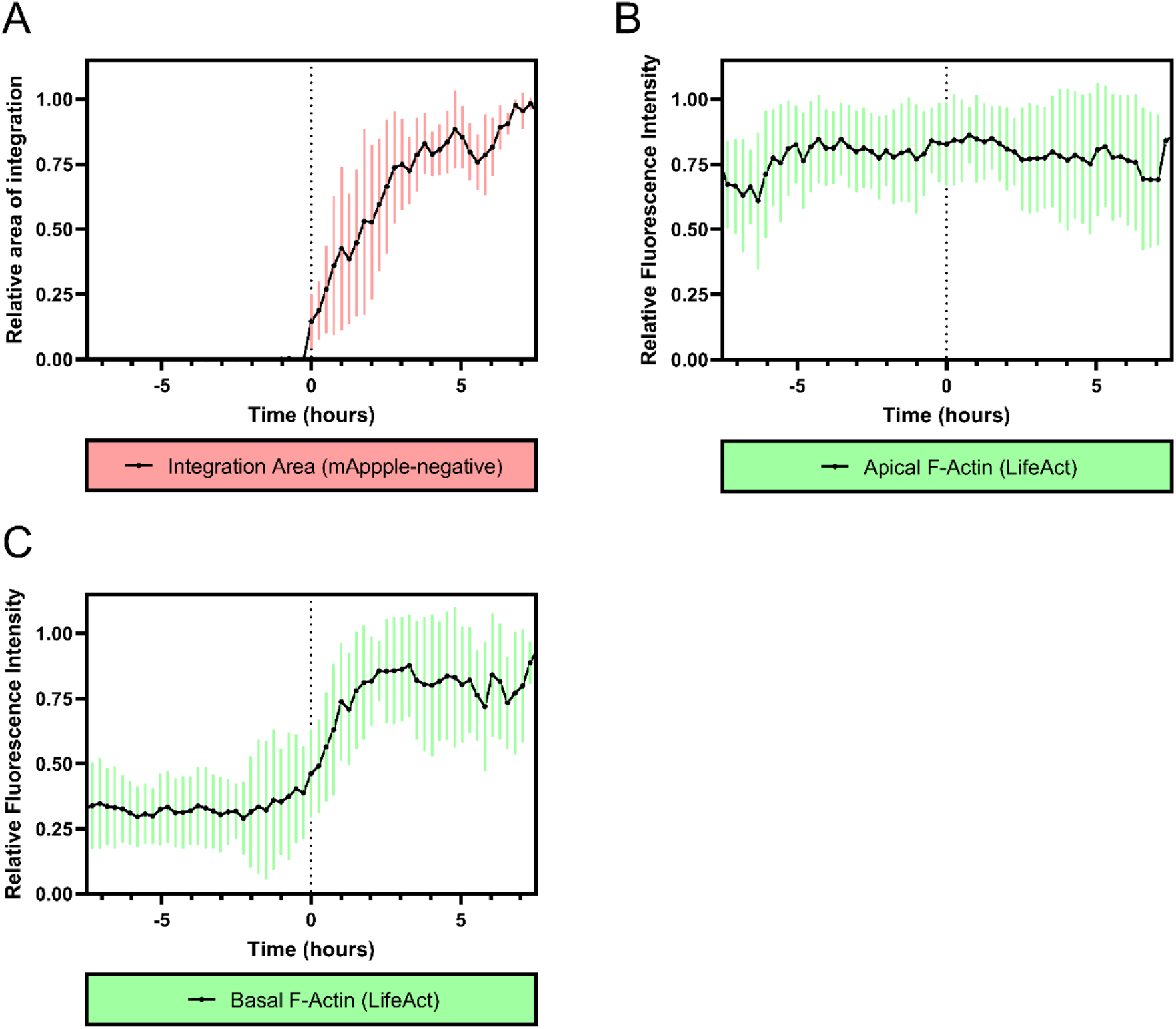
Live imaging of F-actin rearrangement associated with intercalation. A.-C. Quantification of integration area and F-actin intensity at apical interface and basal surface from live imaging analysis of intercalation events. Example shown in Fig. 6 B. Time normalized to initial intercalation denoted at time = 0h. n = 8 cells, Mean +/- 1SD. Fluorescence intensity, area, and time are normalized. A. Area displaced in the monolayer by intercalating cell. B. F-Actin (LifeAct) intensity at apical region (interface) of intercalating cell relative to total LifeAct intensity. C. F-Actin (LifeAct) intensity at basal/bottom surface of intercalating cell relative to total LifeAct intensity.

**Supplemental video 1 & 2**

Examples from live imaging experiments showing cells intercalating into an existing monolayer. Monolayer is labeled with mApple in red channel and cells atop the monolayer can be seen in contrast in the DIC channel.

**Supplemental video 3 – 5**

Examples of live imaging experiments showing cells intercalating (LifeAct-GFP+) into an existing monolayer (mApple+). X,Y axis shown in to and X,Z axis (orthogonal slice) shown below. Line denotes plane in Z or Y axis shown in the other view.

